# Cryo-EM Sample Preparation with Soft-Landing and Laser Flash Melting

**DOI:** 10.1101/2025.06.05.657968

**Authors:** Sarah V. Barrass, Tim K. Esser, Nathan J. Mowry, Lukas Eriksson, Jakub Hruby, Laurence T. Seeley, Wyatt A. Curtis, Lindsay A. Baker, Marcel Drabbels, Stephan Rauschenbach, Ulrich J. Lorenz

**Affiliations:** Ecole Polytechnique Fédérale de Lausanne (EPFL), Laboratory of Molecular Nanodynamics, CH- 1015 Lausanne, Switzerland; Kavli Institute for Nanoscience Discovery, Dorothy Crowfoot Hodgkin Building, University of Oxford, South Parks Road, Oxford OX1 3QU, UK; Department of Chemistry, University of Oxford, Mansfield Road, Oxford OX1 3TA, UK; Department of Biochemistry, University of Oxford, South Parks Road, Oxford OX1 3QU, UK

## Abstract

The preparation of cryo-EM samples by soft-landing mass spectrometry promises to significantly simplify sample optimization, which has remained an important bottleneck in single-particle cryo-electron microscopy (cryo-EM). However, only compacted configurations are observed after soft-landing since the proteins are dehydrated in the process. Here, we demonstrate that proteins can be returned to their native state by depositing a layer of amorphous ice and briefly flash melting the sample to rehydrate the soft-landed proteins. Melting with a 30 µs laser pulse creates a broad ensemble of partially rehydrated conformations. However, a subset of particles recovers their native configuration. This allows us to propose strategies to fully rehydrate the entire ensemble and brings routine sample preparation by soft-landing mass spectrometry within reach. Finally, the experiments also expand the toolbox of microsecond time-resolved cryo-EM for initiating a broader range of protein dynamics.

---

Improvements in instrumentation and data analysis are continuing to expand the scope of cryo-EM ^1–12^, which is rapidly becoming the preferred method for determining the structures of biological macromolecules. However, obtaining cryo-EM samples suitable for high-resolution imaging has remained a significant bottleneck ^13–19^. Contamination of the sample by small molecules frequently reduces the contrast, while the presence of damaged particles or other proteins can make it challenging to obtain high-resolution reconstructions. Samples prepared by plunge freezing may also suffer from an uneven protein distribution across the specimen support as well as variations in ice thickness, with too few areas thin enough for high-resolution imaging ^17^. Preparing cryo-EM samples by native soft-landing electrospray ion beam deposition (ESIBD) potentially presents an appealing alternative that should be able to overcome these issues ^20–24^. Here, proteins are transferred into the gas-phase via electrospray ionization, mass selected, and gently deposited onto a specimen support held at cryogenic temperature. This offers the advantage that a mass filter can be used to select the protein of interest, while removing contaminations and broken particles. Moreover, soft-landing generates a uniform protein distribution across the specimen support. Deposition of a layer of amorphous ice after soft-landing protects the proteins during imaging and improves the attainable spatial resolution. This has recently made it possible to obtain a reconstruction of soft-landed β-galactosidase at 2.6 Å ^25^. However, the proteins are almost entirely dehydrated as they are transferred into the gas phase, so that the soft-landed particles do not present the native protein structure. While secondary structural elements are preserved, water-filled spaces within the proteins usually contract, giving the structures a compacted appearance ^26^.

Here, we demonstrate that soft-landed proteins can be returned to their native, solution-phase structure by briefly flash melting the amorphous ice sample with a laser beam. We have previously shown that flash melting and revitrifying cryo-EM samples preserves the structure of the proteins, allowing for near-atomic resolution reconstructions from revitrified samples. Moreover, the process can be employed to perform microsecond time-resolved observations of protein dynamics ^27–34^, an approach that we extend here to rehydrate soft-landed proteins. In a recent publication, this method has been applied to β-galactosidase particles landed from a molecular ion beam ^35^. The experiments showed a change in appearance of the particles, but yielded only a low-resolution reconstruction, making it difficult to assess how the resulting protein structure is related to the native state. Here, we provide a detailed analysis of the rehydration process, showing that flash melting with a 30 µs laser pulse produces a broad ensemble of partially rehydrated conformations. Crucially, a subset of the population recovers its native structure, which paves the way for using soft-landing mass spectrometry as tool for preparing cryo-EM samples.

Figure 1 illustrates the experimental concept. Samples of soft-landed proteins are prepared with a modified mass spectrometer that is equipped with a cryogenic deposition stage (Supplementary Information A,B) ^25,36^. Proteins (pink) are transferred from solution into the gas phase via native electrospray (Fig. 1A) and enter the vacuum system of the mass spectrometer. During this process, the proteins are dehydrated (blue). The protein ions are mass selected (Fig. 1B) and thermalized in a collision cell to narrow their energy distribution. Finally, they are gently deposited at a collision energy of less than 2 eV per charge onto a 2 nm thick amorphous carbon film on a holey gold specimen support (1.2 µm holes, 1.3 µm apart on a 300-mesh gold grid) held at 110 K (Fig. 1C). To rehydrate the soft-landed proteins, the sample is transferred to a transmission electron microscope modified for time-resolved experiments (Supplementary Information C) ^37,38^. A 50 nm thick layer of amorphous ice is deposited *in-situ* by leaking water vapor into the volume surrounding the sample, which is held at a temperature of 100 K (Fig. 1D). The sample is then flash melted with a 30 µs laser pulse (Fig. 1E, 532 nm wavelength), allowing the proteins to rehydrate (Fig. 1F). Once the heating laser is switched off, the sample rapidly cools and revitrifies within microseconds ^27^, trapping the proteins in their new configurations (Fig. 1G). Finally, the sample is transferred to a high-resolution cryo microscope for imaging.

**Figure 1.**
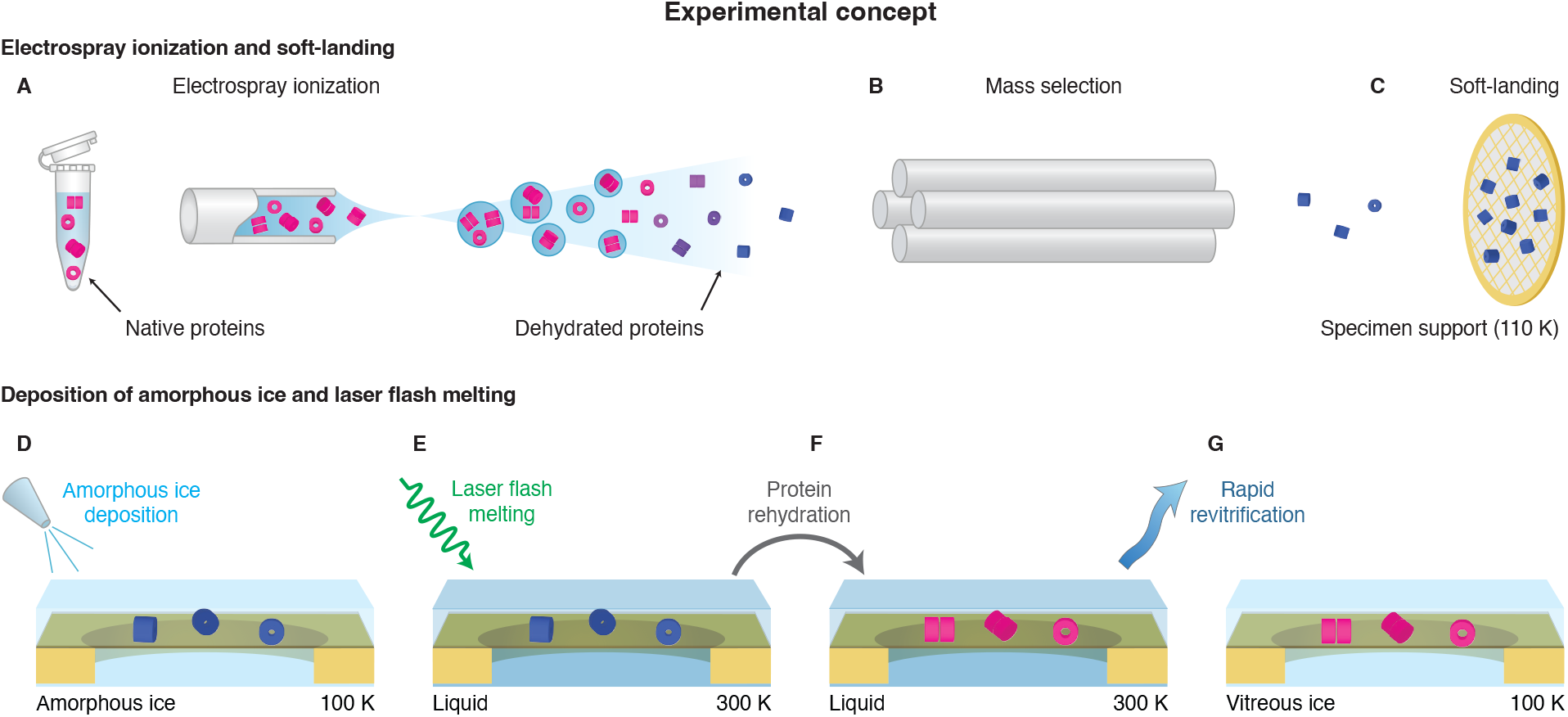
Experimental concept. (**A–C**) Native soft-landing electrospray ion beam deposition (ESIBD). Protein ions are generated by electrospray ionization (**A**), mass selected (**B**), and soft-landed onto an amorphous carbon specimen support held at cryogenic temperature (**C**). In the process, the ions are dehydrated. (**D–H**) Rehydration through deposition of amorphous ice and laser flash melting. Amorphous ice is deposited (**D**) and the sample is laser flash melted (**E**) to all allow the proteins to rehydrate (**F**). Once the heating laser is switched off, the sample rapidly cools and revitrifies (**G**).

Figure 2 compares reconstructions of soft-landed and rehydrated GroEL with a reconstruction from a conventional, plunge-frozen sample (Supplementary Information D–F). While the soft-landed particles only yield a low-resolution reconstruction (Fig. 2A, nominal resolution of 6.8 Å), depositing an additional layer of amorphous ice (1–2 nm thickness) on the proteins reduces radiation damage during imaging and improves the resolution (Fig. 2B, 4.8 Å). A comparison with the native structure (Fig. 2D, 1.9 Å resolution) reveals that dehydration has significantly compacted the soft-landed particles, reducing their height and diameter by about 24 % and 7 %, respectively. The water-filled spaces between the subunits that are present in the native state have contracted, with only the central cavity of the protein still visible. In contrast, flash melting and revitrifying the sample with a 30 µs laser pulse allows the soft-landed particles to rehydrate and expand, restoring the water-filled spaces between the subunits and resulting in a map with dimensions close to that of the native structure (Fig. 2C). The resolution of 3.9 Å is lower than in a conventional, plunge-frozen sample. This suggests that the rehydrated ensemble has significant conformational heterogeneity and that during the time window provided by the 30 µs melting laser pulse, most particles have not fully rehydrated. This is supported by a joint variability analysis of the rehydrated and native particles, which shows that their distributions are separated, with only some overlap at the wings (Fig. S9).

**Figure 2.**
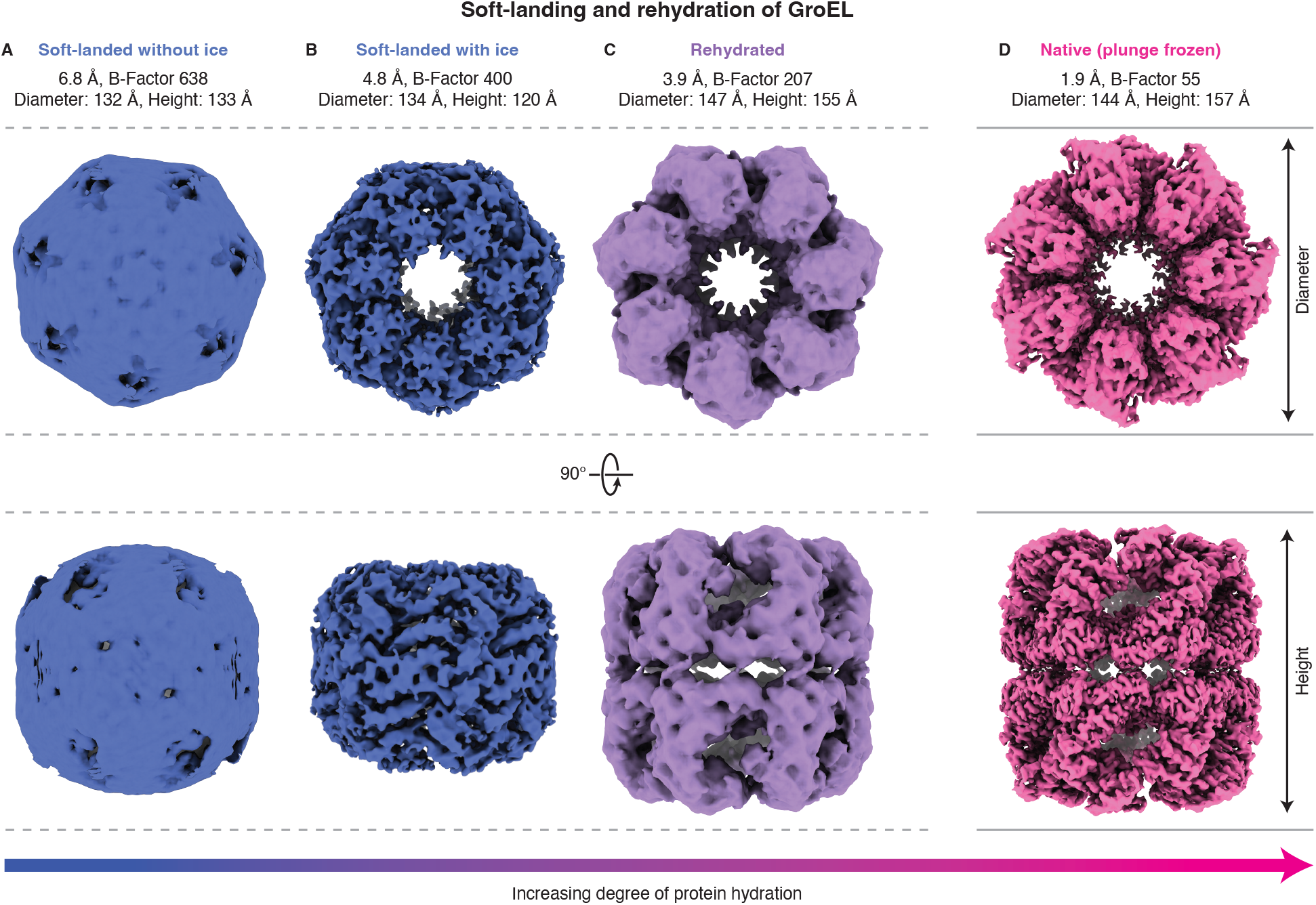
Single-particle reconstructions of soft-landed, rehydrated, and native GroEL. (**A–D**) Maps of GroEL (D_7_ symmetry) after soft-landing (**A**), embedded in amorphous ice (**B**), and after laser melting and rehydration (**C**) are compared with a map of native GroEL from a plunge-frozen sample (**D**). Note that the nominal resolution of the reconstruction in (**A**) does not seem to accurately describe the true resolution.

A soft-landing and rehydration experiment with β-galactosidase ^25^ provides a more detailed understanding of the rehydration process and reveals that laser melting is able to return a subset of the particles to their native state (Fig. 3, Supplementary Information D–F). Similarly to GroEL, the bare, soft-landed β-galactosidase particles only yield a low-resolution reconstruction (Fig. 3A). While depositing amorphous ice improves the resolution (Fig. 3B), it is not able to return the particles to their native structure (Fig. 3E). In contrast, flash melting with a 30 µs laser pulse allows the proteins to rehydrate and expand (Fig. 3C). However, the resulting map appears slightly twisted compared to the native state, with parts of the protein surface exhibiting a low resolution (Fig. S7D). Variability analysis ^39^ of the combined rehydrated and native particles confirms that laser flash melting has not allowed most particles to fully return to their native structure (Supplementary Information G). Figure 3F displays the particle distributions along the first variability component, which is largely associated with the expansion of the particles as they rehydrate. The ensemble of the rehydrated particles (purple) is separated from the native particles (pink), with only the tail of the distribution overlapping with the native ensemble. If we select the particles from this tail, which are most similar to the native state (about 25’000, marked with a dashed box), we obtain the reconstruction in Fig. 3D. Within the resolution obtained (3.7 Å), the structure is indistinguishable from that of native β-galactosidase, demonstrating that during the short time window provided by the melting laser pulse, a subset of the particles return to their native structure. This is also reflected in the similarity of the molecular models that we have built for both structures (Supplementary Information H, Fig. S12, root-mean-square deviation of 0.7 Å), as illustrated in Fig. 3G–I for the subunit contacts within the core of the protein. Note that the reconstruction in Fig. 3D has a worse resolution in parts of the protein surface (Fig. S8E), indicating larger conformational heterogeneity than in a conventional cryo-EM sample, where the particles are fully equilibrated. This is consistent with the B-factor of 62 being larger than for a reconstruction from a conventional sample (B-factor of 42). For comparison, the reconstruction from all rehydrated particles has a B-factor of 103 (a reconstruction with C_1_ symmetry of the particles most similar to the native state has a B-factor of 41, Fig. S11).

**Figure 3.**
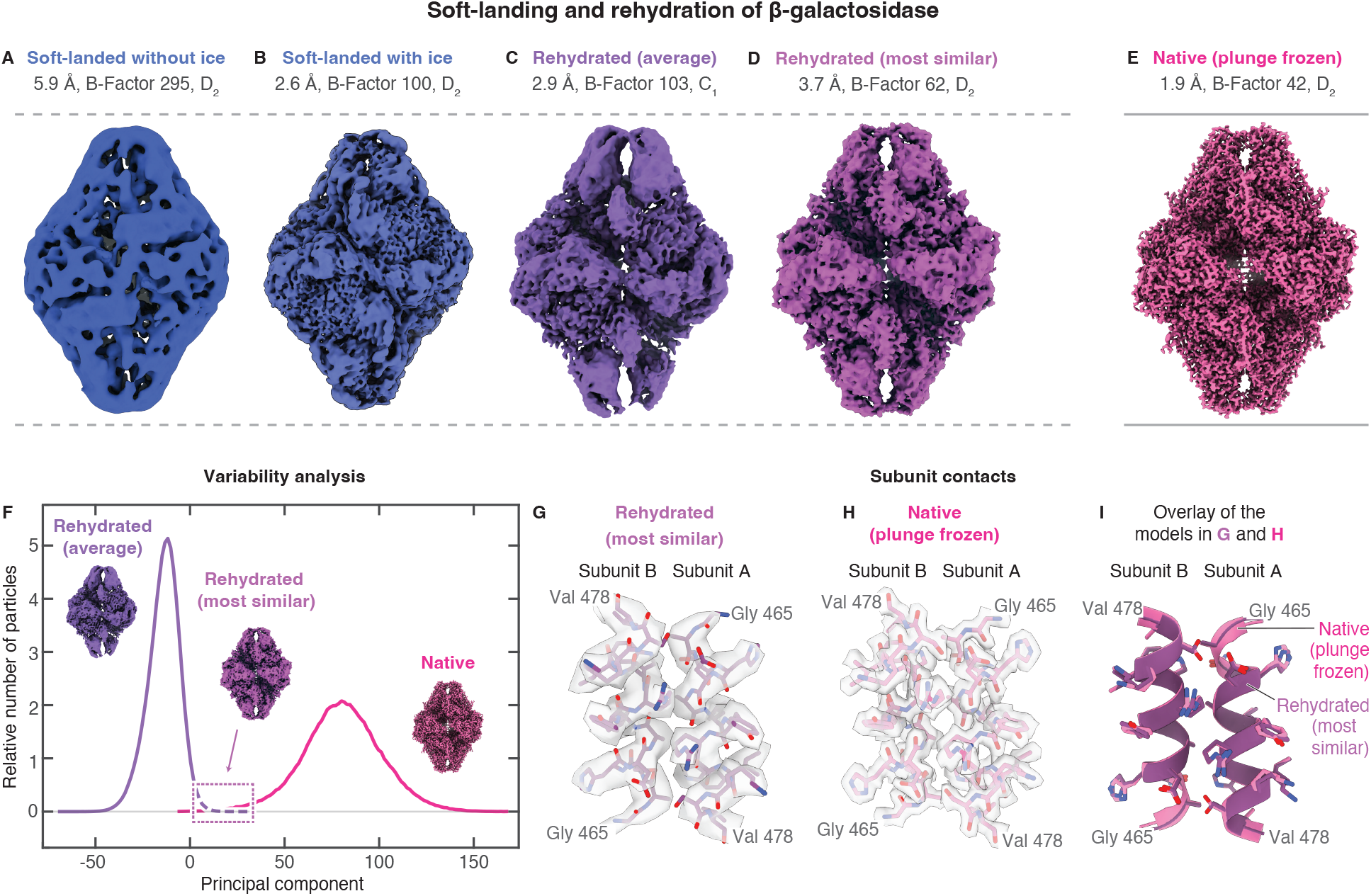
Single-particle reconstructions of soft-landed, rehydrated, and native β-galactosidase. (**A–E**) Maps of β-galactosidase after soft-landing (**A**), embedded in amorphous ice (**B**, EMD-18244 ^25^), and the average structure obtained after laser melting and rehydration (**C**) are compared with a map of the native structure from a plunge-frozen sample (**E**). The volume in (**D**) represents a reconstruction of the rehydrated particles that most closely resemble the native state, as determined from the variability in (**F**). All reconstructions are performed with D_2_ symmetry, with the exception of the map in (**C**), for which no symmetry was applied. The resolution, B-factor, and symmetry of each reconstruction are indicated. (**F**) Joint variability analysis of the rehydrated (purple) and native particles (pink). The particle distributions are shown as a function of the first principal component. A box marks the subset of rehydrated particles that were used for the reconstruction in (**D**). (**G–I**) Comparison of the subunit contacts in the maps of the most similar rehydrated particles (**G**) and native particles (**H**), displayed at 4 σ and 8 σ above the mean, respectively. The corresponding molecular models closely resemble each other as revealed by an overlay in (**I**).

A detailed analysis of the conformational ensemble of the rehydrated particles suggests that adsorption at an interface slows the rehydration process, causing the particles to adopt asymmetric configurations. When the bare, soft-landed particles (Fig. 4A) are rehydrated, their faces expand, with a gap between the subunits widening, as seen in the average structure of the rehydrated particles (Fig. 4B, highlighted with arrows). Close inspection reveals that this expansion is asymmetric, with one face of the structure more expanded than the other. We find that this asymmetry decreases along the first principal component of the variability analysis of Fig. 3F, with the particles becoming more symmetric the closer they are to the native state (Fig. S13). The rehydrated particles most similar to the native structure barely show any asymmetry (Fig. 4C, see Fig. S11 for a reconstruction with C_1_ symmetry). Remarkably, the decrease in asymmetry along the principal component is accompanied by a decrease in preferred orientation of the particles. This can be seen if we divide the central portion of the distribution of the rehydrated particles into four subsets as illustrated in Fig. 4D and determine the angular distribution for each (Fig. 4F–I, shown for one of two samples from which data were recorded, as detailed in Fig. S13). The particles in subset 1, which are the most different from the native structure and the least symmetric, exhibit the most pronounced preferred orientation (Fig. 4F), with one particle view predominantly populated that corresponds to the viewing direction in Fig. 4B (left).

**Figure 4.**
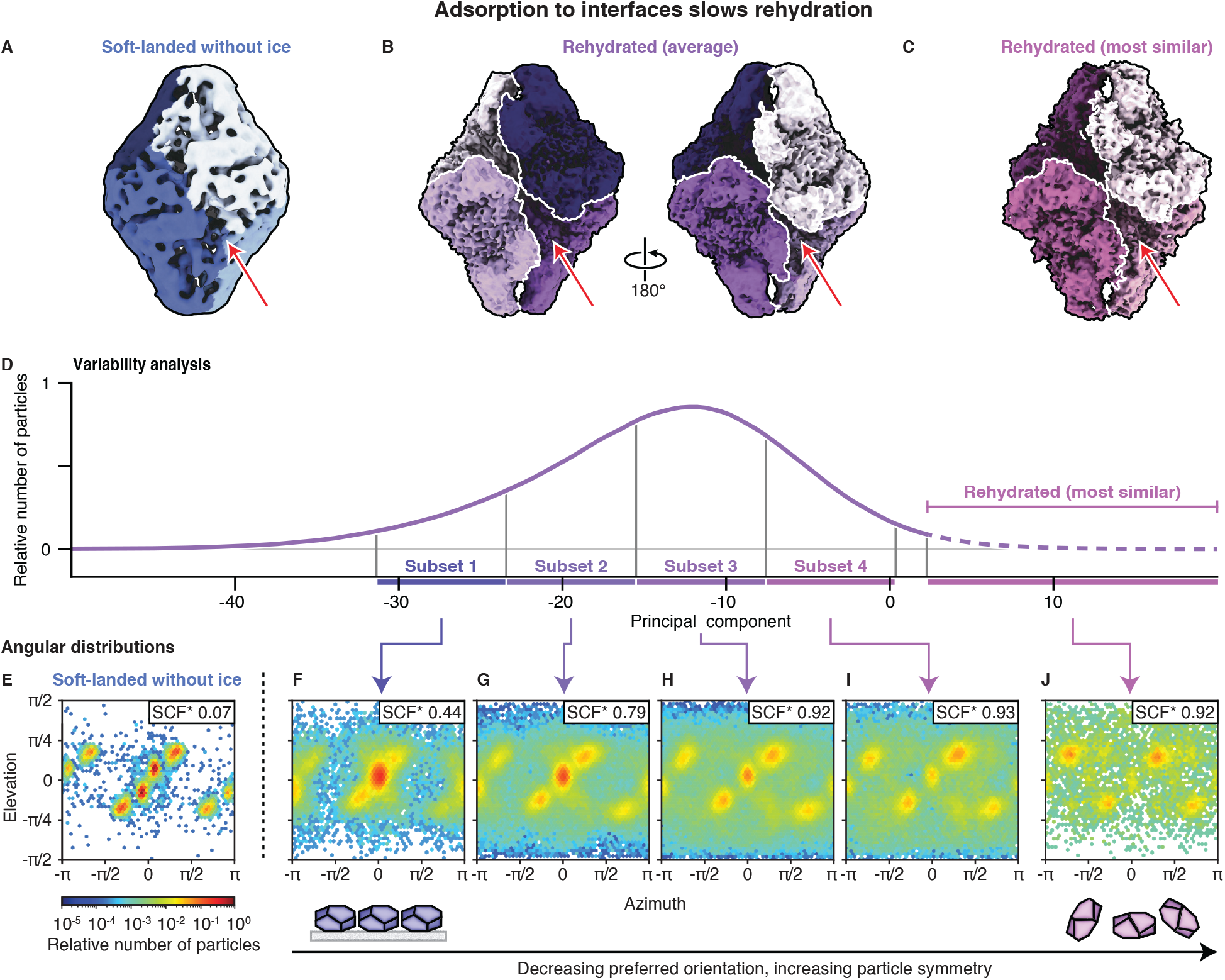
Adsorption to interfaces slows rehydration and causes the particles to adopt asymmetric configurations. (**A–C**) Reconstructions of β-galactosidase for the bare, soft-landed particles (**A**, from Fig. 3A), all rehydrated particles (**B**, from Fig. 3C), and the rehydrated particles most similar to the native structure (**C**, from Fig. 3D). The subunits are highlighted in colors. The average structure of the rehydrated particles in (**B**) shows significant asymmetry in the expansion of its two faces (arrows). (**D**) The central part of the distribution of rehydrated particles along the variability component in Fig. 2F is divided into four subsets. The particles included in the reconstruction in (**C**) are also indicated. (**E–J**) Angular distributions of the bare, soft-landed particles (**E**), of the four subsets of rehydrated particles (**F–I**), and of the rehydrated particles most similar to the native state (**J**). The angular distributions of the rehydrated particles are shown for one of two samples, with the other presented in Fig. S13. The sampling compensation factor (SCF*), a measure of the degree of preferred orientation, is indicated. Along the principal component, preferred particle orientation decreases, and the particles become more symmetric (Fig. S13), suggesting that adsorption causes the particles to rehydrate asymmetrically.

Preferred orientation results from the adsorption of particles to an interface of the cryo-EM sample ^40–42^. Evidently, most particles in subset 1 are adsorbed with one of their faces, which strongly suggests that this slows the expansion of this face, breaking the symmetry of the particles as they rehydrate. Along the principal component, particles adsorbed in this orientation become less frequent (Fig. 4G–I). Moreover, the angular distribution turns more homogeneous, with the sampling compensation factor (SCF*), a measure of the degree of preferred orientation, improving from 0.44 in subset 1 to 0.93 in subset 4. The SCF* takes values between zero and one, with one corresponding to a perfectly isotropic angular distribution. The rehydrated particles that are most similar to the native state and most symmetric show little preferred orientation, with an SCF* of 0.92 (Fig. 4J). Their angular distribution shows maxima that are only sparsely populated in subset 1, suggesting that adsorption in these orientations allows the particles to rehydrate more easily.

Our results suggest that adsorption to an interface hinders many particles from fully returning to the native state during the short time window provided by the melting laser pulses. It is conceivable that some particles simply remain adsorbed to the amorphous carbon film of the specimen support onto which they were soft-landed. It is also possible that once the sample is liquid, other particles quickly diffuse to the air-water interface (estimated diffusion time on the order of 10 µs) and adsorb there. The angular distribution of the most asymmetric particles suggests that both scenarios may occur. It features a second, less pronounced maximum that corresponds to the opposite viewing direction (Figs. 4F and S13C), suggesting that particles also adsorb to the opposite sample interface. It is difficult to decide which interface the particles predominantly adsorb to merely based on the preferred orientation of the most asymmetric particles, since it resembles both that of the bare, soft-landed particles (Fig. 4E) and that of the particles in a conventional cryo-EM sample, where adsorption to the air-water interface is likely dominant (Fig. S5F). It should however be possible to distinguish between both scenarios by taking care to mount the sample in a well-defined orientation during image acquisition, which we have not done here.

In conclusion, our experiments outline a path towards preparing cryo-EM samples with soft-landing mass spectrometry by depositing amorphous ice onto the soft-landed particles and using laser flash melting to rehydrate them. This approach allows us to prepare samples with a uniform particle distribution across the specimen support. While a conventional sample of β-galactosidase contains some fraction of dimers (Fig. S5C), only tetramers are found after soft-landing (Figs. S6,7), underlining the potential of this approach to reduce sample heterogeneity. We observe that during the time window provided by a 30 µs melting laser pulse, most particles only partially rehydrate, which results in a broad conformational distribution. However, for β-galactosidase, a subset of particles does return to its native state. This suggests that it should ultimately be possible to fully rehydrate all particles simply by providing them with more time to do so.

We have recently demonstrated that the time window during which the laser pulse keeps the sample liquid can be extended from tens to hundreds of microseconds by enclosing the cryo-EM sample between two ultrathin silicon dioxide layers ^33^. These sealing membranes, which we vapor deposit onto the sample, stabilize the thin liquid film under laser irradiation and prevent its evaporation. For compact proteins such as β-galactosidase, this should provide enough time for most particles to fully rehydrate. Our experiments also suggest, that the rehydration process can be accelerated if adsorption of the particles to interfaces can be reduced. As we have recently shown, the hydrophilic silicon dioxide membranes eliminate interactions of the particles with the hydrophobic air-water interface ^33^. It should also be possible to minimize interactions with the support film by optimizing the choice of material or by depositing a thin layer of amorphous ice prior to soft-landing. Future experiments will also have to show whether this approach can be generalized to a wide range of proteins. For example, some particles require the presence of specific ions or buffer components to maintain their native fold. For such proteins, it should be possible to co-deposit these compounds in the desired stoichiometry.

Finally, our experiments also suggest a new approach for initiating dynamics in microsecond time resolved cryo-EM experiments ^34^. It should be possible to use soft-landing mass spectrometry to deposit a ligand or small molecule onto a cryo-EM sample, so that upon laser flash melting, rapid mixing occurs, and the compound can interact with the embedded proteins to initiate dynamics. This provides the advantage that one would not be limited to the availability of suitable photocaged compounds in order to initiate dynamics and should open up new possibilities for studying signaling, drug binding, or even protein-protein interactions.

## Data availability

The cryo-EM maps have been deposited in the Electron Microscopy Data Bank (EMDB) under accession codes EMD-52627 (soft-landed GroEL without ice), EMD-52626 (soft-landed GroEL with ice), EMD-52284 (soft-landed and rehydrated GroEL), EMD-52282 (plunge-frozen GroEL), EMD-52274 (soft-landed β-galactosidase without ice), EMD-52244 (soft-landed and rehydrated β-galactosidase (average)), EMD-52260 (soft-landed and rehydrated β-galactosidase (most similar)), EMD-52230 (plunge-frozen β-galactosidase). The corresponding data set are accessible on EMPIAR (EMPIAR-12558, EMPIAR-12576, EMPIAR-12481, EMPIAR-12480, EMPIAR-12489, EMPIAR-12479, EMPIAR-12475). The atomic coordinates of the models have been deposited in the Protein Data Bank (PDB) under accession codes 9HKI (plunge-frozen β-galactosidase) and 9HLM (soft-landed and rehydrated β-galactosidase (most similar)).

## Acknowledgements

We acknowledge the support of the Dubochet Center for Imaging (DCI) in Lausanne, and the COSMIC microscopy facility in Oxford. This work was supported by the ERC Starting Grant 759145 (awarded to U.J.L.), the Swiss National Science Foundation Grant PP00P2_163681 (awarded to U.J.L.), the UK Research and Innovation (UKRI) Grant EP/V051474/1 (awarded to S.R.), the H2020 Marie Curie Actions Grant 883387 (awarded to T.K.E.), and the Wellcome Trust Grant 218482/Z/19/Z (awarded to L.T.S.).

## Author contributions

T.K.E. conceptualized this work. L.T.S. performed the plunge freezing. T.K.E. conducted solution preparation and soft-landing. N.J.M. and S.V.B. conducted the sample rehydration. S.V.B., J.H., and L.E. performed the data acquisition. S.V.B and L.E. performed the cryo-EM data processing. S.V.B. performed the structure modelling and refinement. S.V.B. and W.A.C. analyzed the data. S.R., M.D., and U.J.L. supervised the work. The writing of the original draft was performed by S.V.B. and T.K.E. The data visualization was performed by S.V.B., J.H, and W.A.C. The reviewing and editing of the manuscript were performed by all authors.

