## Supplementary Information for "Cryo-EM Sample Preparation with Soft-Landing and Laser Flash Melting"

**Affiliations:**

† Current affiliation: Thermo Fisher Scientific, De Schakel 2, 5651GH Eindhoven, Noord-Brabant, The Netherlands

‡ Current affiliation: Ecole Polytechnique Fédérale de Lausanne (EPFL), Laboratory for Ultrafast X-ray Sciences, CH-1015 Lausanne, Switzerland

**This PDF file includes:**

A. Preparation of protein solutions

B. Native mass spectrometry and ion beam deposition onto specimen supports

C. Laser melting and revitrification experiments

D. Preparation of conventional cryo-EM samples

E. Data collection

F. Single-particle reconstructions

G. Variability analysis of soft-landed and rehydrated β-galactosidase

H. Model building

I. Supplementary figures

J. Supplementary tables

**A. Preparation of protein solutions**

Samples of GroEL and β-galactosidase were purchased from Sigma-Aldrich (G3153-5MG and C7688-1MG). For the preparation of conventional cryo-EM samples, solutions were prepared as follows. The lyophilized GroEL powder was resuspended in 20 mM Tris (pH 7.9), 50 mM KCl, 0.5 mM EDTA, 5 mM $MgCl_2$, and 0.5 mg/mL ATP to give a protein concentration of 4 mg/mL. A solution of β-galactosidase (1.35 mg/mL) was prepared by resuspending in 25 mM Tris (pH 7.9), 50 mM NaCl, 10 mM EDTA, and 2 mM $MgCl_2$. For native mass spectrometry, the GroEL solution was desalted by passing it through two P6 buffer exchange columns (7326222, Biorad) that had been equilibrated with 200 mM ammonium acetate (pH 6.9). A solution of β-galactosidase (10 mM) was obtained by dissolving the lyophilized powder in 200 mM ammonium acetate and desalting with the same procedure.

**B. Native mass spectrometry and ion beam deposition onto specimen supports**

Native mass spectrometry was performed on a commercial mass spectrometer (Thermo Scientific Q Exactive UHMR instrument) that was modified for ion beam deposition experiments [1–3]. The following operating parameters were typically used.

| | |
|---|---|
| Source DC offset | 21 V |
| S-lens RF level | 200 (300 $V_{p\text{-}p}$) |
| Transfer capillary temperature | 60 °C |
| Ion transmission settings | “high m/z” (700 $V_{p\text{-}p}$ for the injection flatapole and 900 $V_{p\text{-}p}$ for the bent flatapole, transfer multipole, and collision cell) |
| Detector optimization | “high m/z” |
| Injection flatapole | 5 V |
| Interflatapole lens | 4 V |
| Bent flatapole | 2 V |
| Transfer multipole | 0 V |
| Collision-cell pressure setting | 7 ($N_2$) |
| Collision-cell multipole DC | −5 V |
| Collision-cell exit lens | −15 V |

Nano-electrospray ionization emitters were fabricated by pulling borosilicate glass capillaries (30-0042, Harvard Bioscience) with a pipette puller (P-1000, Sutter Instrument) and gold coating them with a sputter coater (108A/SE, Cressington). Up to 10 µl of protein solution was loaded into an emitter, and the emitter tip was clipped to create an opening of 0.1 to 10 µm [4]. Electrospray ionization was initiated by applying a potential of 1.2 kV and a gentle nanoflow pressure (<200 mbar above atmospheric pressure).

While the conventional operating mode of the mass spectrometer was used to collect mass spectra [5], a modified scan matrix was employed for ion deposition that allows the ions to pass through the C-trap and collision cell without being trapped and reach the deposition stage. The number of water molecules attached to the protein complex ions under these conditions is estimated to be less than 200, as determined by comparing with mass spectra recorded after desolvating the ions in the collision cell. Note that the protein complex ions may also contain residual salt.

Protein complex ions were soft-landed on specimen supports as previously described [1–3]. Clipped UltrAuFoil specimen grids covered with a 2 nm thick layer of amorphous carbon (R1.2/1.3, 300 mesh, Quantifoil) were mounted in a shuttle, which was inserted into the deposition stage of the modified mass spectrometer[3] a few minutes before the experiment. The deposition stage is cooled to a temperature of 110 K by a closed-cycle helium cryo-cooler (Sunpower Cryotel GTLT). When the shuttle is inserted, a spring-loaded mechanism opens a cover, exposing the grid for the ion beam deposition. At the same time, an electrical contact is created, which is used to control the sample potential and with it the deposition energy. This contact is also used to monitor the ion deposition current with a picoammeter, so that the protein coverage on the specimen grid can be controlled. Protein ions were deposited with an energy of 2 eV per charge, as measured with a retarding-grid energy detector, with a typical ion current of 50 pA and 20 min deposition time. The deposition is performed in ultrahigh vacuum ($< 10^{-9}$ mbar).

For the reconstructions in Figs. 2B and 3B (GroEL and β-galactosidase, “soft-landed with ice”), a <10 nm thick layer of amorphous ice was additionally deposited onto the specimen grids after ion beam deposition. To this end, the partial pressure of water vapor in the deposition chamber was raised to $5 \cdot 10^{-5}$ mbar using a gas dosing valve, while the sample temperature was maintained between 111 K and 114 K. After a typical deposition time of about 1 min, the partial pressure of water was reduced by more than 2 orders of magnitude, and the sample was cooled to about 95 K to prevent devitrification during sample transfer. Finally, the grids were transferred into liquid nitrogen in under 2 min, as previously described [3]. The deposition rate was calibrated separately by depositing layers of amorphous ice under identical conditions and determining their thicknesses with electron tomography [6].

**C. Laser melting and revitrification experiments**

Laser melting and revitrification experiments were performed with a modified JEOL 2010F transmission electron microscope [7,8]. Microsecond laser pulses for flash melting (30 µs, 532 nm wavelength, 83–149 mW laser power) were generated by chopping the output of a continuous laser (Coherent Verdi G7) with an acousto-optic modulator (AA Opto-Electronic) controlled by an arbitrary function generator (Tektronix AFG1062). The laser beam was focused to a spot size of 38 µm FWHM on the sample, as determined from an image of the laser beam recorded with a CCD camera placed in a conjugate plane. The laser beam was centered onto a grid square in order to melt and revitrify a sample area.

Samples prepared by soft-landing (no additional ice coverage) were loaded into the microscope with a Simple Origin Dual Autogrid cryo specimen holder, 100 K sample temperature. Amorphous ice was then deposited onto the sample *in situ* by leaking water vapor into the area surrounding the sample with a gas dosing valve as previously described [7,8]. With the shutter of the cryo specimen holder still covering the sample to prevent any deposition of amorphous ice, the gas dosing valve was opened, and the deposition rate was set to about 15 nm/min (calibrated in a separate experiment as described before [7,8]). After the deposition rate had been stable for 10 min, the shutter was opened to deposit a 50 nm thick layer of amorphous ice (25 nm on each side). The deposition was stopped by covering the sample again with the shutter. The gas dosing valve was then closed, and at least one hour was allowed for the column of the microscope to return to its base pressure, before the shutter was opened again to perform the laser revitrification experiments.

**D. Preparation of conventional cryo-EM samples**

Conventional cryo-EM samples of GroEL and β-galactosidase were prepared after centrifuging the protein solutions to remove aggregates (15 min, 18000 g, 4 ºC). For GroEL, 3 µL of the protein solution was applied to holey amorphous carbon grids (R2/1, 200 mesh copper, Quantifoil) that had been rendered hydrophilic through glow-discharging. The grids were blotted for 5s and plunged into a liquid ethane/propane mixture using a manual plunger. Cryo-EM samples of β-galactosidase were prepared with the same procedure, using a volume of 4 µL, holey carbon grids covered with a 2 nm thick amorphous carbon layer (R2/1, 400 mesh copper, Quantifoil), and liquid ethane/propane as a cryogen.

**E. Data collection**

Micrographs of soft-landed GroEL with and without an additional ice layer deposited on the sample (Fig. 2A,B) were collected at the COSMIC Cryo-EM Facility, University of Oxford, UK using a Talos Arctica 200 kV microscope equipped with a Falcon 4i detector (without ice) and with a Titan Krios G3 equipped with a GIF Bioquantum energy fiter and a Gatan K3 camera (with ice). All other datasets were collected at the Dubochet Center for Imaging in Lausanne, Switzerland, using a Titan Krios G4 microscope equipped with a Falcon 4i detector and a SelectrisX energy filter. The data acquisition parameters for the different data sets are listed in Tables S1 and S2.

**F. Single-particle reconstructions**

<u>Plunge-frozen GroEL</u>

Single-particle reconstructions were performed in CryoSPARC v4.5. The processing workflow is shown in Fig. S1. The micrographs were subjected to patch motion-correction, and movies with a total full-frame motion distance of more than 50 px were discarded. CTF estimation was performed using Patch CTF, and micrographs with an estimated resolution of worse than 6 Å, astigmatism greater than 1000 Å, or relative ice thickness values above 1.5 or below 0.9 were discarded. Particles were picked using a blob picker with a diameter of 120-180 Å, extracted with a box size of 512 px, and subjected to two rounds of 2D classification. The best classes were selected and used for template-based picking using a particle diameter of 150 Å. The particles were extracted with a box size of 512 px. After one round of 2D classifications, the selected particles were used for *ab-initio* reconstruction with two classes, resulting in one class representative of the intact protein (Fig. S4, blue), and one junk class (Fig. S4, grey). Following heterogeneous refinement, non-uniform refinement ($D_7$ symmetry) was performed on the intact protein class, with local and global CTF refinement and Ewald sphere correction, yielded a map at a resolution of 1.9 Å. The global and local resolution were estimated using gold-standard FSC thresholds of 0.143 and 0.5, respectively. The apical region of the map exhibits a lower resolution, which 3D classification (four classes) followed by non-uniform refinement of each class ($C_7$ symmetry), revealed to be caused by the presence of different states. Docking molecular models into the volumes allowed us to identify an ATP-bound and an ADP-bound state (PDB: 4AAQ[9] and PDB: 4KI8[10], respectively), which are shown in Fig. S10. The structures were visualized with UCSF ChimeraX 1.7.

Soft-landed GroEL without ice

Single-particle reconstructions were performed in CryoSPARC 4.1. The processing workflow is shown in Fig. S2. The micrographs were subjected to patch motion-correction, and movies with a total full-frame motion distance of more than 50 px were discarded. CTF estimation was performed using Patch CTF and micrographs with a defocus tilt angle of greater than 15.5 degrees discarded. Particles were manually picked from one micrograph, extracted with a box size of 352 px, and subjected to three rounds of 2D classification. The best classes were selected and used for template-based picking using a particle diameter of 150 Å. The particles were extracted with a box size of 300 px, subjected to three rounds of 2D classification, followed by *ab-initio* reconstruction with one class. The particles were refined using heterogeneous refinement with three classes. Particles from classes two and three were then refined using non-uniform refinement with $D_7$ symmetry, against an initial model generated by low pass filtering the published cryo-EM map EMD-8750 to 30 Å. This yielded a map at a resolution of 6.7 Å. The global and local resolution were estimated using gold-standard FSC thresholds of 0.143 and 0.5, respectively. The structures were visualized with UCSF ChimeraX 1.7.

Soft-landed GroEL with ice

Single-particle reconstructions were performed in CryoSPARC 4.1. The processing workflow is shown in Fig. S3. The micrographs were subjected to patch motion-correction. CTF estimation was performed using Patch CTF and micrographs with an estimated resolution of worse than 8.13 Å, average intensities above 507.24 or below -407.35, or relative ice thickness values above 1.19 were discarded. Particles were picked using template-based picking with templates generated from manual picking and a particle diameter of 140 Å. The particles were extracted with a box size of 352 px and subjected to three rounds of 2D classification. The best classes were selected for *ab-initio* reconstruction with five classes, followed by heterogeneous refinement with five classes. The best class was then refined with homogeneous refinement followed by non-uniform refinement ($D_7$ symmetry). This yielded a map at a resolution of 4.8 Å. The global and local resolution were estimated using gold-standard FSC thresholds of 0.143 and 0.5, respectively. The structures were visualized with UCSF ChimeraX 1.7.

<u>Soft-landed and rehydrated GroEL</u>

Single-particle reconstructions were performed in CryoSPARC v4.5. The processing workflow is shown in Fig. S4. Micrographs were collected from two grids. The micrographs were subjected to patch motion-correction, and movies with a total full-frame motion distance of more than 50 px were discarded. CTF estimation was performed using Patch CTF, and micrographs with an estimated resolution of worse than 6 Å, or relative ice thickness values above 1.2 or below 1.0 were discarded. In addition, micrographs identified as containing hexagonal ice, cubic ice or no ice by manual inspection were discarded. Templates were generated from the plunge-frozen GroEL reconstruction, low pass filtered to 20 Å and used for template-based picking using a particle diameter of 150 Å. The particles were extracted with a box size of 512 px. Particles extracted from grid one (1,722,594 particles) were subjected to two rounds of 2D classification, and the selected particles were used for *ab-initio* reconstruction, followed by non-uniform refinement ($D_7$ symmetry). Next, all template-picked particles from both grids were classified in 3D against the refined volume and ten decoy classes (random noise generated from the initial guess in an *ab-initio* reconstruction job) using heterogeneous refinement. Particles from the class corresponding to GroEL were then classified twice in 3D against the refined volume and five decoy classes. Finally, good particles were refined using non-uniform refinement ($D_7$ symmetry), yielded a map at a resolution of 3.9 Å. The global and local resolution were estimated using gold-standard FSC thresholds of 0.143 and 0.5, respectively. The structures were visualized with UCSF ChimeraX 1.7.

Plunge-frozen β-galactosidase

Single-particle reconstructions were performed in CryoSPARC v4.5 [11]. The processing workflow is shown in Fig. S5. The micrographs were subjected to patch motion-correction, and movies with a total full-frame motion distance of more than 50 px were discarded. Contrast transfer function (CTF) estimation was performed using Patch CTF, and micrographs with an estimated resolution of worse than 6 Å, astigmatism greater than 1000 Å, or relative ice thickness values above 1.5 or below 0.9 were discarded. Particles were picked using a blob picker with a diameter of 80-180 Å, extracted with a box size of 512 px, and subjected to one round of 2D classification. The best classes were selected and used for template-based picking using a particle diameter of 180 Å. Following template picking, the particles were extracted with a box size of 512 px. After two rounds of 2D classifications, the selected particles were used for *ab-initio* reconstruction with two classes, resulting in one class representative of the intact tetrameric protein (Fig. S1, blue), and one class of the dimeric protein (Fig. S1, grey). Following heterogeneous refinement, and non-uniform refinement ($C_1$ symmetry) of the intact tetrameric protein, 3D classification was performed to remove damaged particles (Fig. S1, grey). Finally, non-uniform refinement ($D_2$ symmetry) with local and global CTF refinement and Ewald sphere correction yielded a map at a resolution of 1.9 Å. The map was post processed using local filtering. The global and local resolution were estimated using gold-standard FSC thresholds of 0.143 and 0.5, respectively. The structures were visualized with UCSF ChimeraX 1.7 [12,13].

Soft-landed β-galactosidase without ice

Single-particle reconstructions were performed in CryoSPARC v4.5 [11]. The processing workflow is shown in Fig. S6. The micrographs were subjected to patch motion-correction, and CTF estimation was performed using Patch CTF. Particles were picked using a blob picker with a diameter of 80-200 Å, extracted with a box size of 540 px, and classified using two rounds of 2D classification. The best classes were selected, and an initial model generated using *ab-initio* reconstruction. The model was refined using non-uniform refinement with $D_2$ symmetry, yielded a map at a resolution of 5.9 Å. The map was post processed using local filtering. The global and local resolution were estimated using gold-standard FSC thresholds of 0.143 and 0.5, respectively. The structures were visualized with UCSF ChimeraX 1.7 [12,13].

Soft-landed and rehydrated β-galactosidase (Average)

Single-particle reconstructions were performed in CryoSPARC v4.5. The processing workflow is shown in Fig. S7. Micrographs were collected from two grids and processed together. The micrographs were subjected to patch motion-correction, and movies with a total full-frame motion distance of more than 50 px were discarded. CTF estimation was performed using Patch CTF, and micrographs with an estimated resolution of worse than 6 Å, or relative ice thickness values above 1.2 or below 1.0 were discarded. In addition, micrographs identified as containing hexagonal ice, cubic ice or no ice by manual inspection were discarded. Particles were picked with a blob picker using a diameter of 80-180 Å, extracted with a box size of 512 px, and subjected to one round of 2D classification. The best classes were selected, and an initial model generated using *ab-initio* reconstruction, before being refined using non-uniform refinement with $D_2$ symmetry. The resulting volume was used to generate templates for template-based picking with a particle diameter of 180 Å. The particles were extracted with a box size of 512 px, subjected to two rounds of 2D classification, followed by *ab-initio* reconstruction with seven classes. The best class was then refined with non-uniform refinement ($C_1$ symmetry). Next, all template-picked particles were classified in 3D against the refined volume and five decoy classes (random noise generated from the initial guess in an *ab-initio* reconstruction job) using heterogeneous refinement, to improve particle classification. Finally, particles were subjected to one round of 2D classification and the good particles refined using non-uniform refinement ($C_1$ symmetry), yielded a map at a resolution of 2.9 Å. The global and local resolution were estimated using gold-standard FSC thresholds of 0.143 and 0.5, respectively. The structures were visualized with UCSF ChimeraX 1.7.

**G. Variability analysis of soft-landed and rehydrated β-galactosidase**

We used variability analysis to understand the motions that the particles undergo as they rehydrate during laser melting. Particles from the soft-landed and rehydrated β-galactosidase dataset (Fig. 3C) were combined with those of the native state (Fig. 3E), and 3D variability analysis in CryoSPARC was performed (filter resolution 10 Å, 3 variability components). Figure 3F shows the conformational distributions along the first component, which predominantly describes the expansion the particles undergo as they rehydrate. The distributions of the rehydrated and the native particles are well separated. This indicates that during flash melting, most particles have not quite approached a fully rehydrated conformation. Only the tail of the distribution overlaps with that of the native particles, suggesting that the particles at this end of the distribution have moved furthest along the trajectory associated with the rehydration. We selected 25,206 particles from this end of the distribution (as indicate in Fig. 3F) and performed an *ab-initio* reconstruction with one class and non-uniform refinement ($D_2$ symmetry) to obtain the map in Fig. 3D, which has a resolution of 3.7 Å (workflow illustrated in Fig. S8).

In order to better understand the rehydration process, we subdivided the distribution of the rehydrated particles along the first component as indicated in Fig. 4D (subsets of even size, with 3 % of the particles left out at either end of the distribution) and generated reconstructions for each subset. A*b-initio* reconstruction with one class and non-uniform refinement ($C_1$ symmetry) yielded the volumes shown in Fig. S11. After aligning the maps, the orientation diagnostics job in cryoSPARC was used to obtain the angular distributions of the particles (Fig. 4F–J).

**H. Model Building**

A model of plunge-frozen β-galactosidase was obtained with the following procedure. An available β-galactosidase model (PDB: 6CVM) was rigidly fit into the locally filtered plunge-frozen β-galactosidase map using UCSF ChimeraX 1.7. The asymmetric unit within the model was then iteratively refined using the real space refine tool in PHENIX 1.21.1-5286-000 [14], with secondary structure and non-crystallographic symmetry constraints imposed, as well as the ISOLDE plugin in UCSF ChimeraX, and Coot [13,15,16].

A model for the soft-landed and revitrified (Best) β-galactosidase map was obtained with a similar procedure. An available β-galactosidase model (PDB: 6CVM [17]) was rigidly fit into the sharpened map using UCSF ChimeraX 1.7. The asymmetric unit was then refined as above while imposing torsional and adaptive distance restraints derived from the starting structure in areas where the backbone density was poorly resolved. Imposing these restraints helped prevent unphysical geometries from occurring during the refinement in the areas of the map with a local resolution between 4 and 7 Å. Finally, the model was trimmed to remove areas where no density was observed. The quality of the models was evaluated using MolProbity [18] and PHENIX.

### I. Supplementary Figures

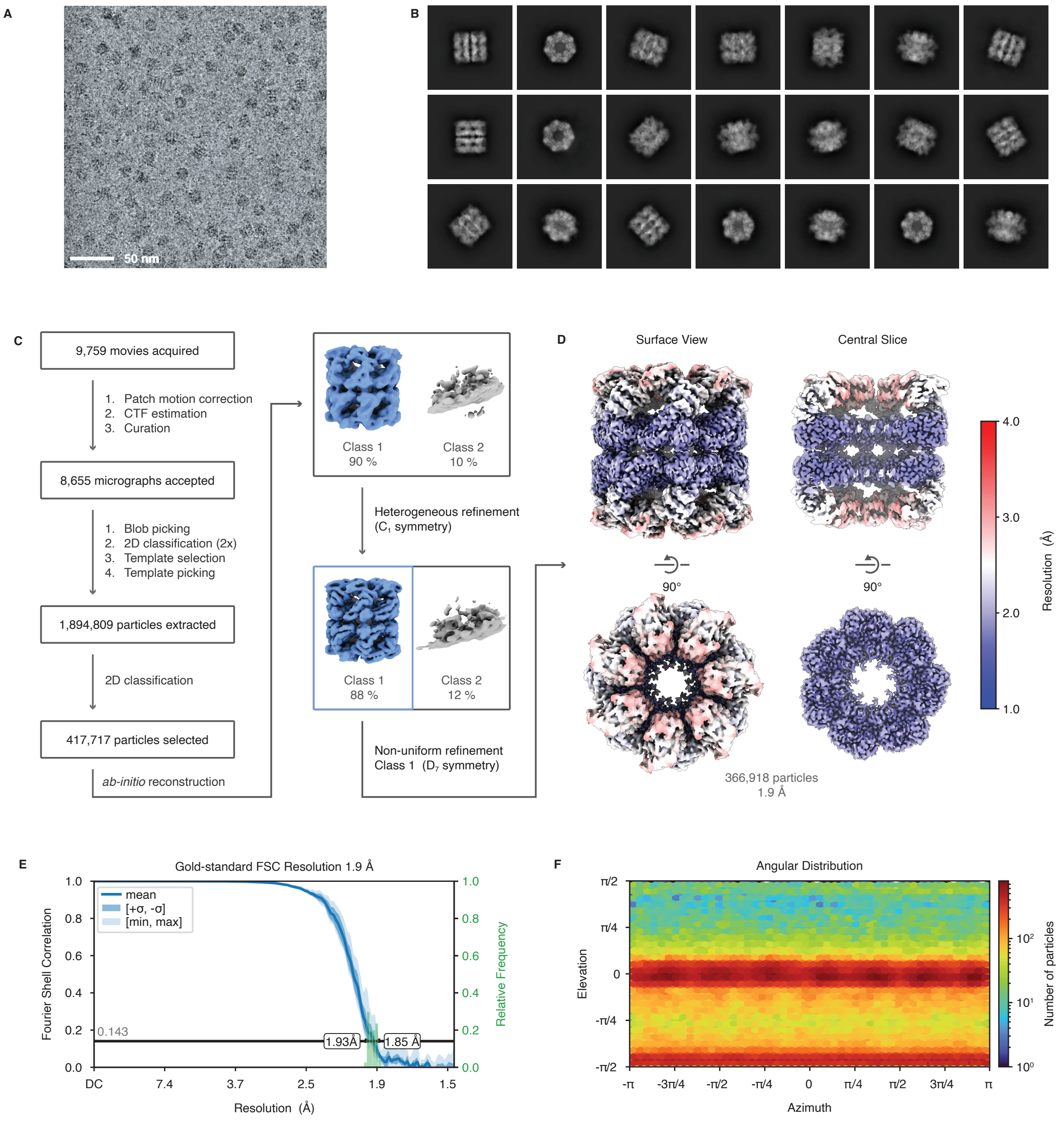


**Figure S1 | Cryo-EM reconstruction of plunge-frozen GroEL.** (**A**) Representative micrograph, low pass filtered by 10 Å. (**B**) Selected 2D class averages of the dataset. (**C**) Single-particle reconstruction workflow in cryoSPARC, yielded a map at a resolution of 1.9 Å (**D**). (**D**) Final 3D reconstruction colored by local resolution. Note that both ADP- and ATP-bound states of GroEL are present in our samples (Fig. S10) [38], which lowers the resolution, particularly in the apical region. (**E**) Conical Gold-Standard Fourier Shell Correlation, with the black line indicating the 0.143 cut-off. The mean value is shown as a solid blue line, with the dark blue shading corresponding to one standard deviation, and the light blue shading indicating the minimum and maximum. A histogram of the resolution values is shown in green; the minimum and maximum resolution are indicated. (**F**) Angular distribution of the particles.

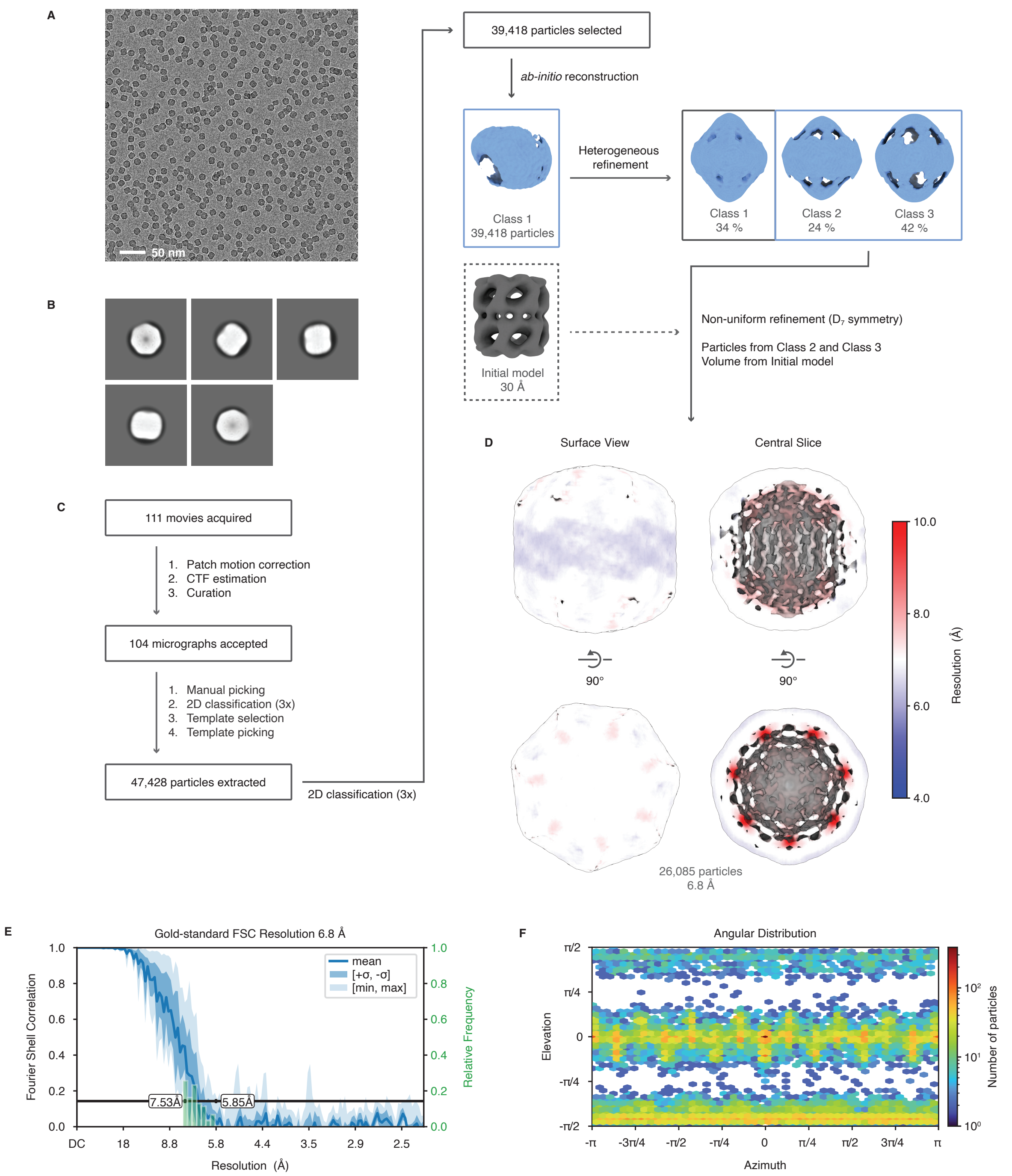


**Figure S2 | Cryo-EM reconstruction of soft-landed GroEL without ice.** (**A**) Representative micrograph. (**B**) Selected 2D class averages of the dataset. (**C**) Single-particle reconstruction workflow in cryoSPARC, yielded a map at a resolution of 6.7 Å (**D**). (**D**) Final 3D reconstruction colored by local resolution. (**E**) Conical Gold-Standard Fourier Shell Correlation, with the black line indicating the 0.143 cut-off. The mean value is shown as a solid blue line, with the dark blue shading corresponding to one standard deviation, and the light blue shading indicating the minimum and maximum. A histogram of the resolution values is shown in green; the minimum and maximum resolution are indicated. (**F**) Angular distribution of the particles.

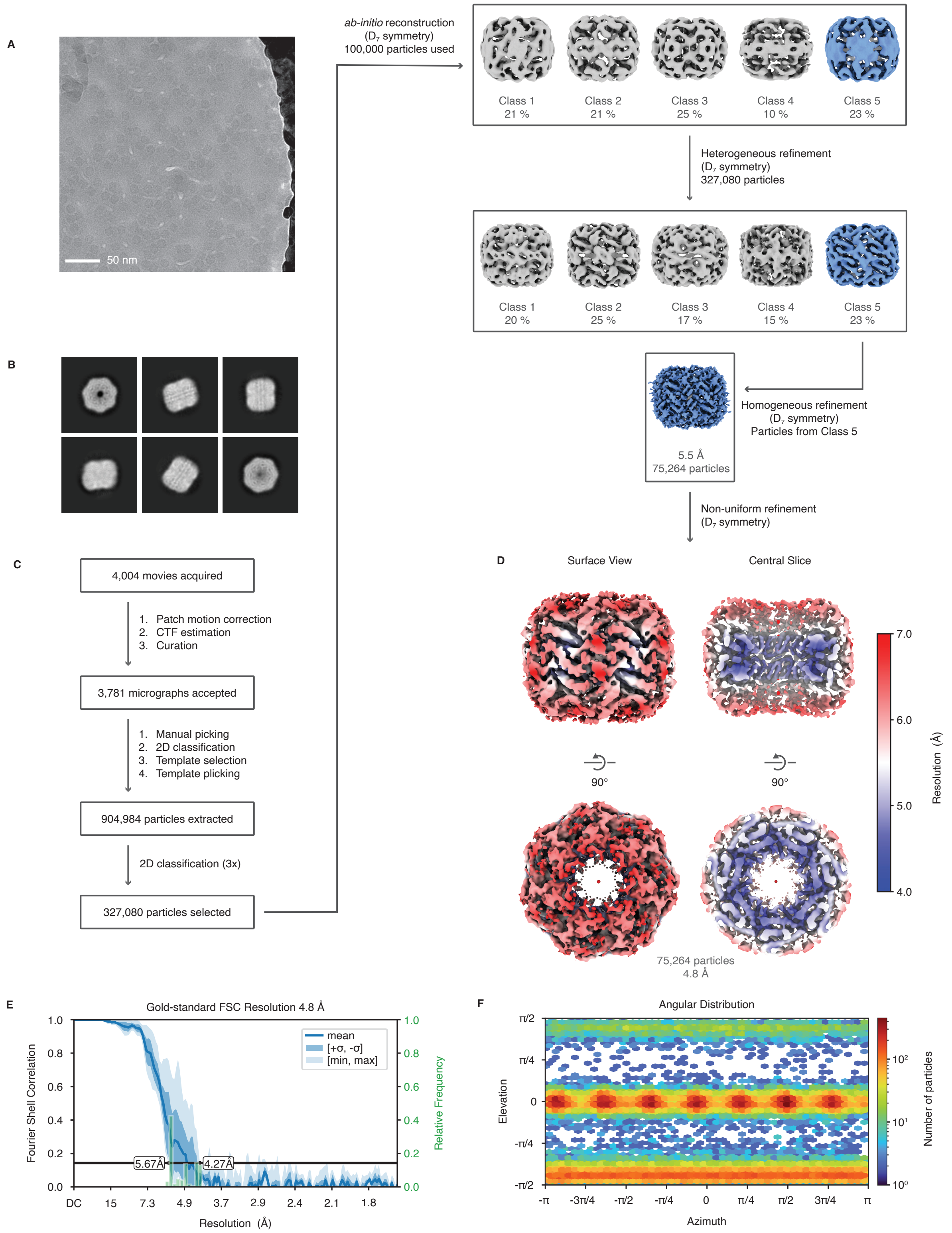


**Figure S3 | Cryo-EM reconstruction of soft-landed GroEL with ice.** (**A**) Representative micrograph. (**B**) Selected 2D class averages of the dataset. (**C**) Single-particle reconstruction workflow in cryoSPARC, yielded a map at a resolution of 4.8 Å (**D**). (**D**) Final 3D reconstruction colored by local resolution. (**E**) Conical Gold-Standard Fourier Shell Correlation, with the black line indicating the 0.143

cut-off. The mean value is shown as a solid blue line, with the dark blue shading corresponding to one standard deviation, and the light blue shading indicating the minimum and maximum. A histogram of the resolution values is shown in green; the minimum and maximum resolution are indicated. (**F**) Angular distribution of the particles.

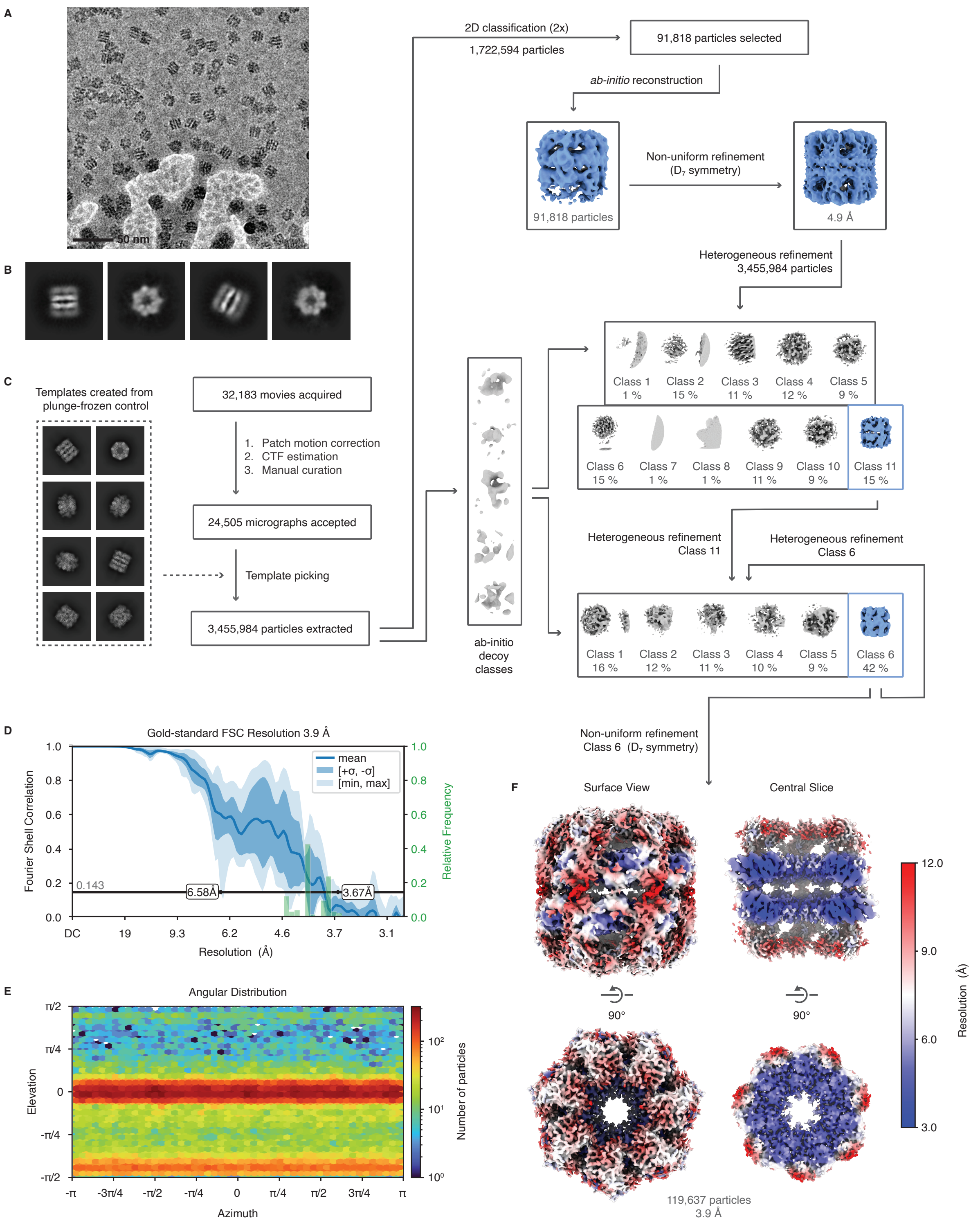


**Figure S4 | Cryo-EM reconstruction of soft-landed and rehydrated GroEL.** (**A**) Representative micrograph, low pass filtered by 10 Å. (**B**) Selected 2D class averages of the dataset. (**C**) Single-particle reconstruction workflow in cryoSPARC, yielded a map at a resolution of 3.9 Å (**F**). (**D**) Conical Gold-Standard Fourier Shell Correlation, with the black line indicating the 0.143 cut-off. The mean value is shown as a solid blue line, with the dark blue shading corresponding to one standard deviation, and the light blue shading indicating the minimum and maximum. A histogram of the resolution values is shown in green; the minimum and maximum resolution are indicated. (**E**) Angular distribution of the particles. (**F**) Final 3D reconstruction coloured by local resolution.

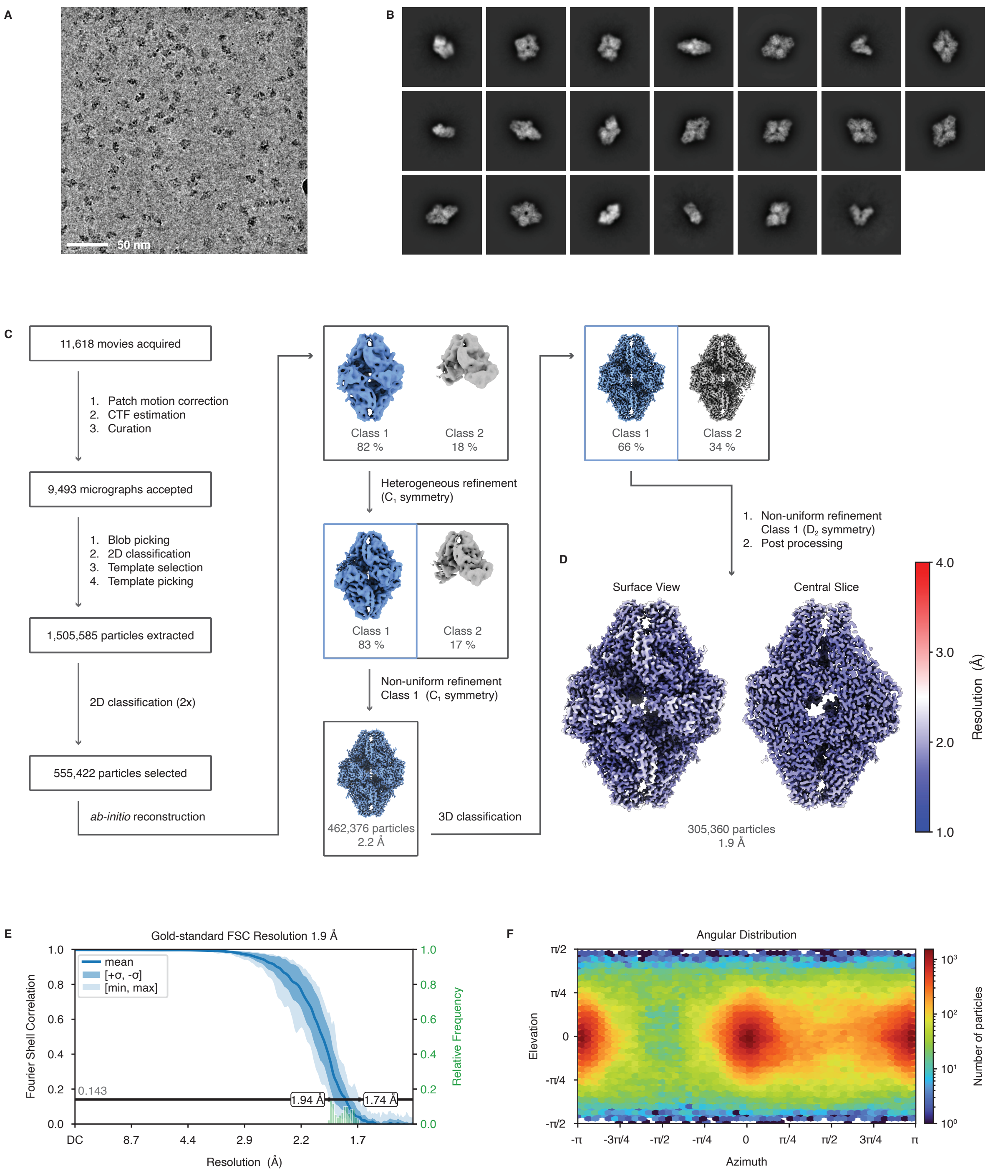


**Figure S5 | Cryo-EM reconstruction workflow for plunge-frozen β-galactosidase.** (**A**) Representative micrograph, low pass filtered by 5 Å. (**B**) Selected 2D class averages of the dataset. (**C**) Single-particle reconstruction workflow in cryoSPARC. (**D**) Final 3D reconstruction with a resolution of 1.9 Å, colored by local resolution. (**E**) Conical Gold-Standard Fourier Shell Correlation, with the black line indicating the 0.143 cut-off. The mean value is shown as a solid blue line, with the dark blue shading corresponding to one standard deviation, and the light blue shading indicating the minimum and maximum. A histogram of the resolution values is shown in green; the minimum and maximum resolution are indicated. (**F**) Angular distribution of the particles.

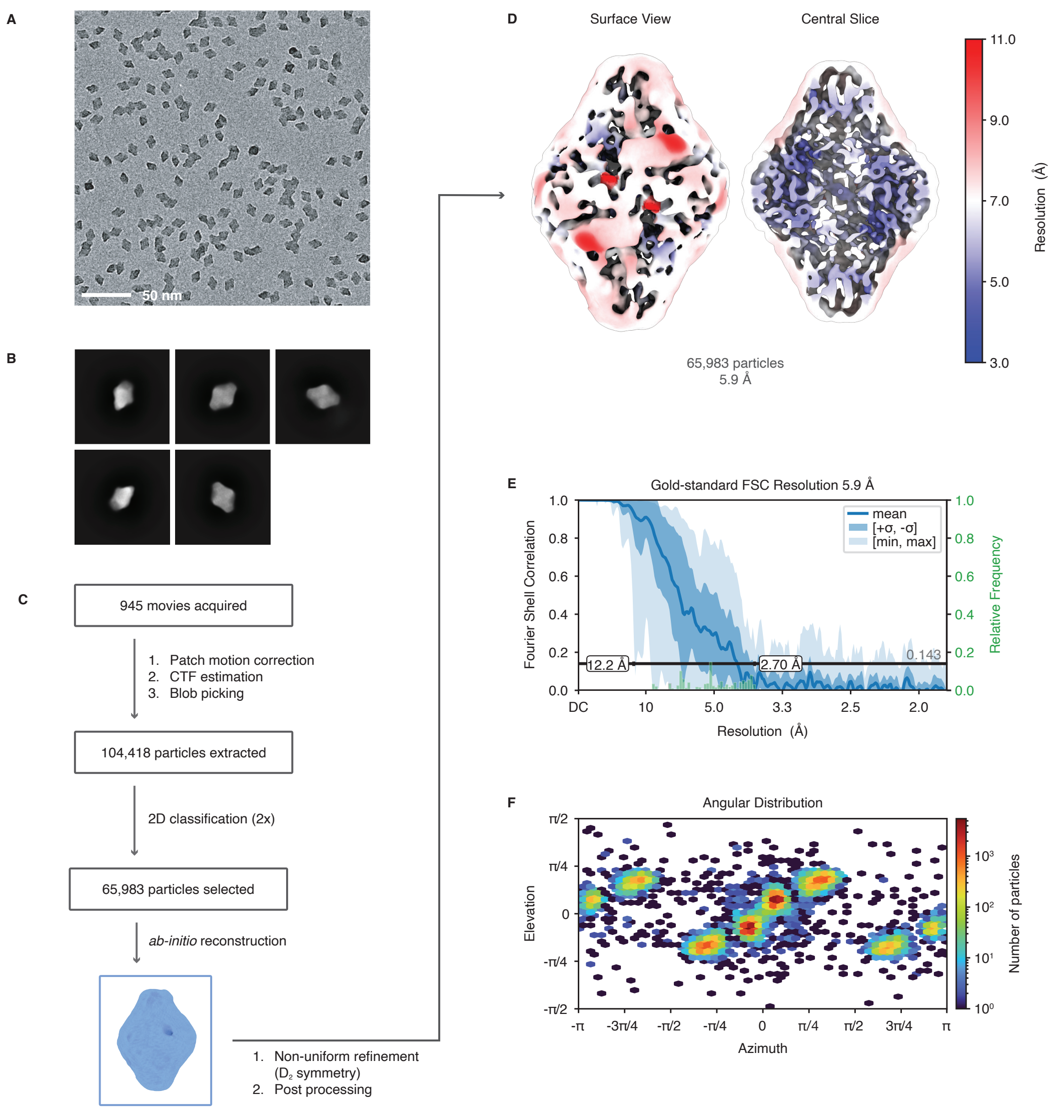


**Figure S6 | Cryo-EM reconstruction workflow for soft-landed β-galactosidase without ice.** (**A**) Representative micrograph. (**B**) Selected 2D class averages of the dataset. (**C**) Single-particle reconstruction workflow in cryoSPARC, yielded a map at a resolution of 5.9 Å (**D**). (**D**) Final 3D reconstruction colored by local resolution. (**E**) Conical Gold-Standard Fourier Shell Correlation, with the black line indicating the 0.143 cut-off. The mean value is shown as a solid blue line, with the dark blue shading corresponding to one standard deviation, and the light blue shading indicating the minimum and maximum. A histogram of the resolution values is shown in green; the minimum and maximum resolution are indicated. (**F**) Angular distribution of the particles.

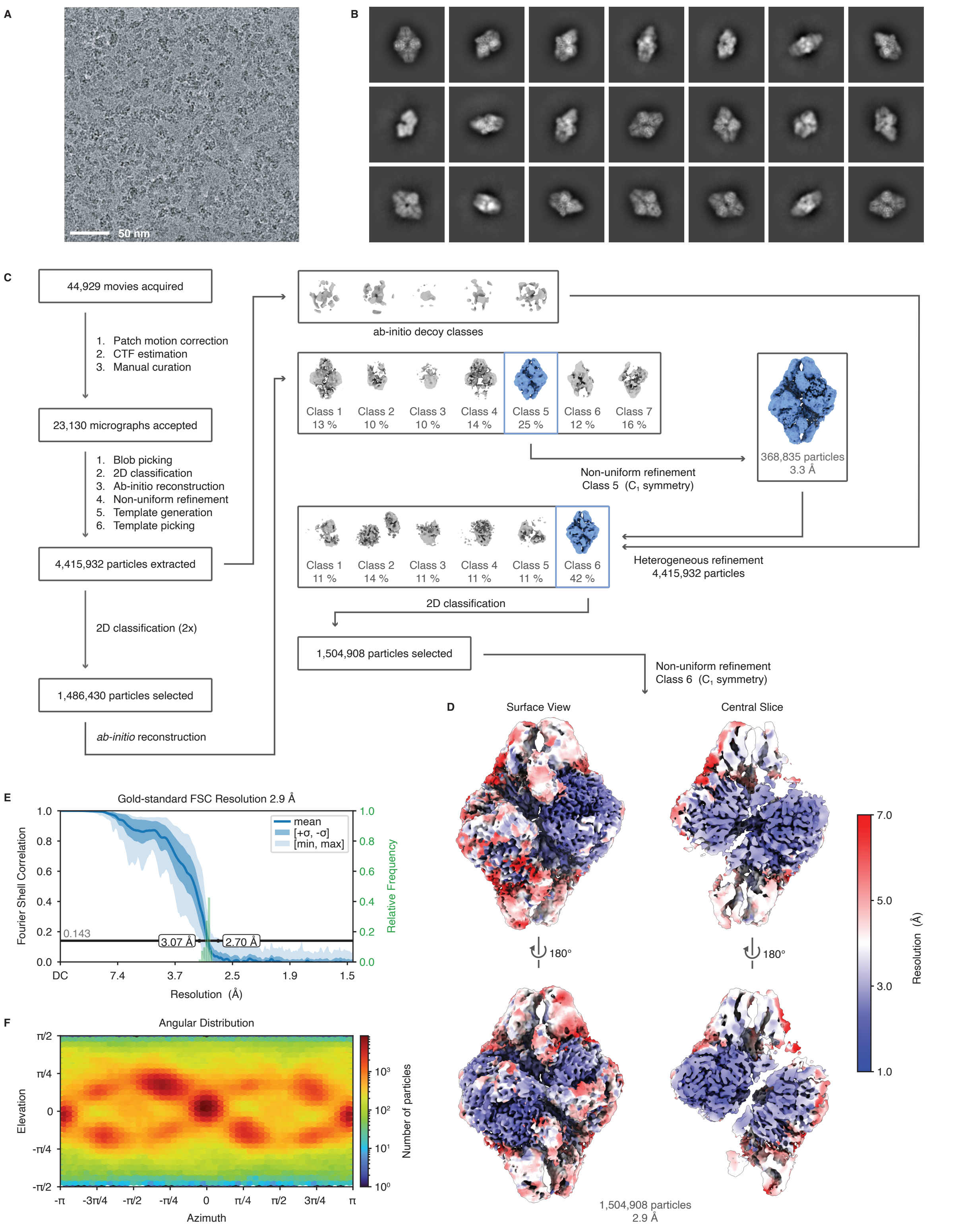


**Figure S7 | Cryo-EM reconstruction workflow for soft-landed and rehydrated β-galactosidase (Average).** (**A**) Representative micrograph, low pass filtered by 5 Å. (**B**) Selected 2D class averages of the dataset. (**C**) Single-particle reconstruction workflow in cryoSPARC, yielded a map at a resolution of 2.9 Å (**D**). (**D**) Final 3D reconstruction colored by local resolution. (**E**) Conical Gold-Standard Fourier

Shell Correlation, with the black line indicating the 0.143 cut-off. The mean value is shown as a solid blue line, with the dark blue shading corresponding to one standard deviation, and the light blue shading indicating the minimum and maximum. A histogram of the resolution values is shown in green; the minimum and maximum resolution are indicated. (**F**) Angular distribution of the particles.

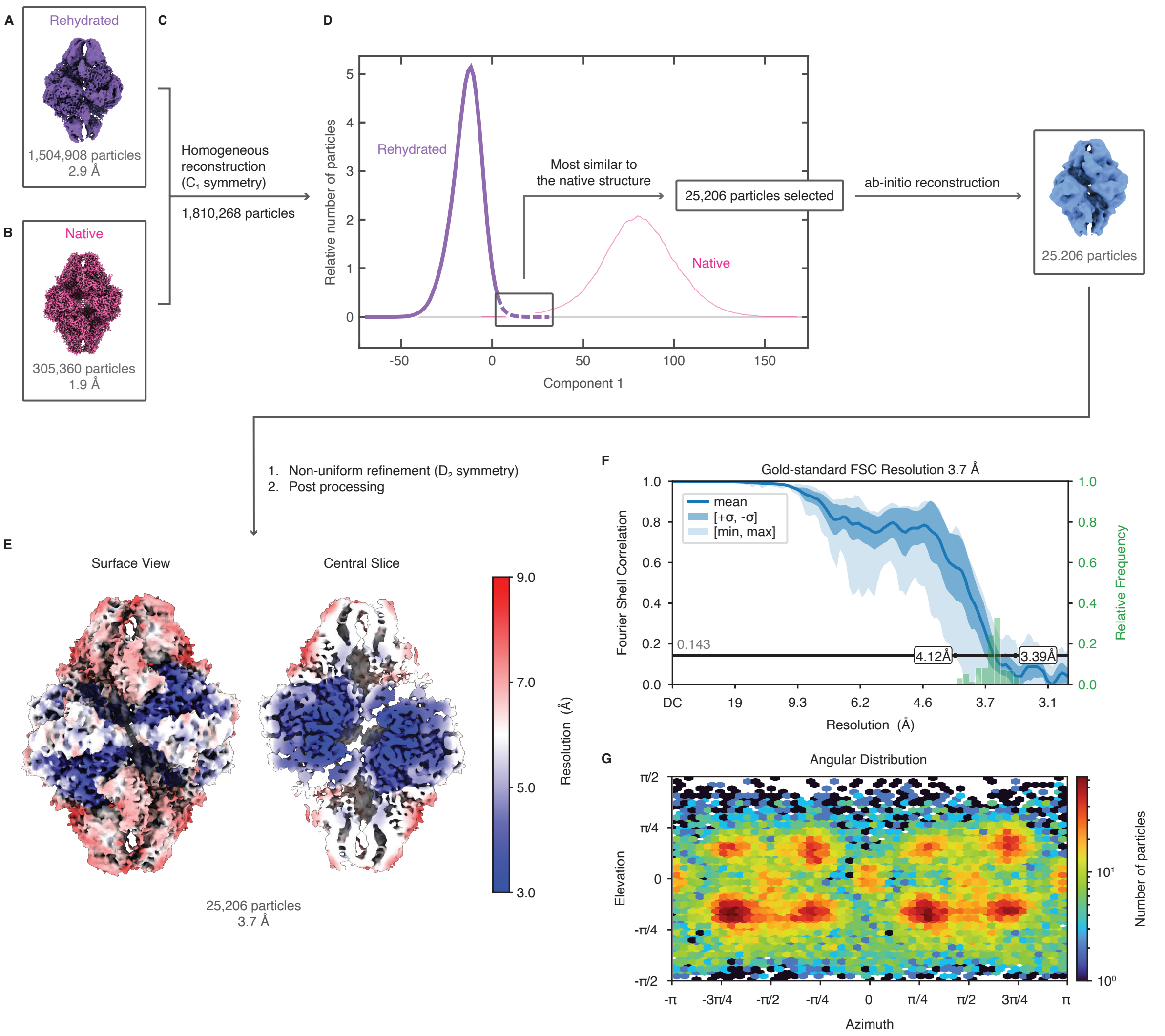


**Figure S8 | Cryo-EM reconstruction workflow for soft-landed and rehydrated β-galactosidase (Best).** (**A**) Reconstruction of soft-landed and rehydrated β-galactosidase (from Fig. 3C). (**B**) Reconstruction of native β-galactosidase from a plunge frozen sample (from Fig. 3E). (**C**) Particles from the reconstructions in (**A**) and (**B**) were combined and subjected to a 3D variability analysis in cryoSPARC. (**D**) Particle distributions along the first variability component, with the rehydrated and native particles shown in purple and pink, respectively. Rehydrated particles from the tail of the distribution (black box) were subjected to *ab-initio* reconstruction and refinement. (**E**) Reconstruction of the rehydrated particles from the tail of the distribution (from Fig. 3D). The map has a resolution of 3.7 Å; the local resolution is indicated in color. (**F**) Conical Gold-Standard Fourier Shell Correlation, with the black line indicating the 0.143 cut-off. The mean value is shown as a solid blue line, with the dark blue shading corresponding to one standard deviation, and the light blue shading indicating the minimum and maximum. A histogram of the resolution values is shown in green; the minimum and maximum resolution are indicated. (**G**) Angular distribution of the particles.

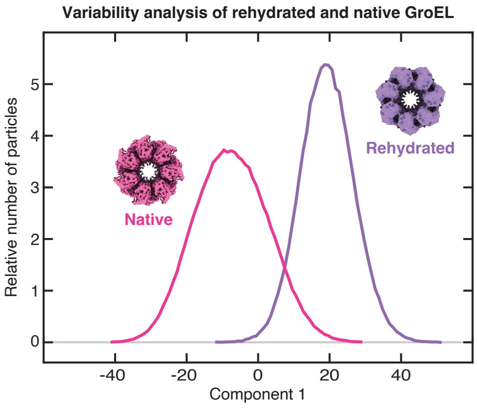


**Figure S9 | Variability analysis of the merged datasets of rehydrated and native GroEL.** The particle distributions are shown as a function of the first variability component, which is mainly associated with the expansion of the particles as they rehydrate. The distributions of the rehydrated and native sample are shown in purple and pink, respectively. The variability analysis was performed in CryoSPARC (filter resolution 7 Å, 3 components).

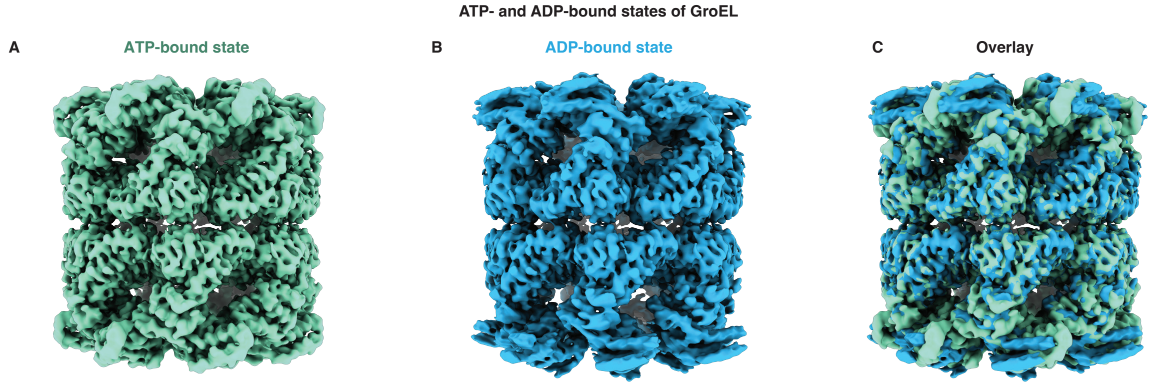


**Figure S10 | Comparison of the ATP- and ADP-bound states of GroEL present in the native (plunge frozen) sample.** (**A**) ATP-bound state. (**B**) ADP-bound state. (**C**) Overlay of (**A**) and (**B**).

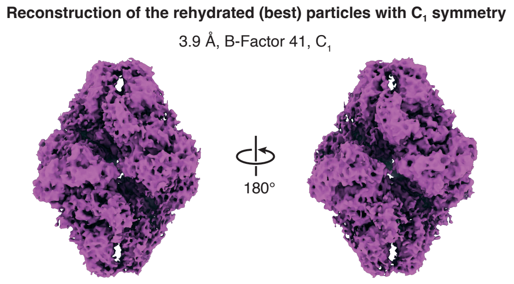


**Figure S11 | Reconstruction of the rehydrated (most similar) particles with $C_1$ symmetry.** Both sides of the particles have expanded symmetrically. The map is shown at a threshold of 5 σ above the mean.

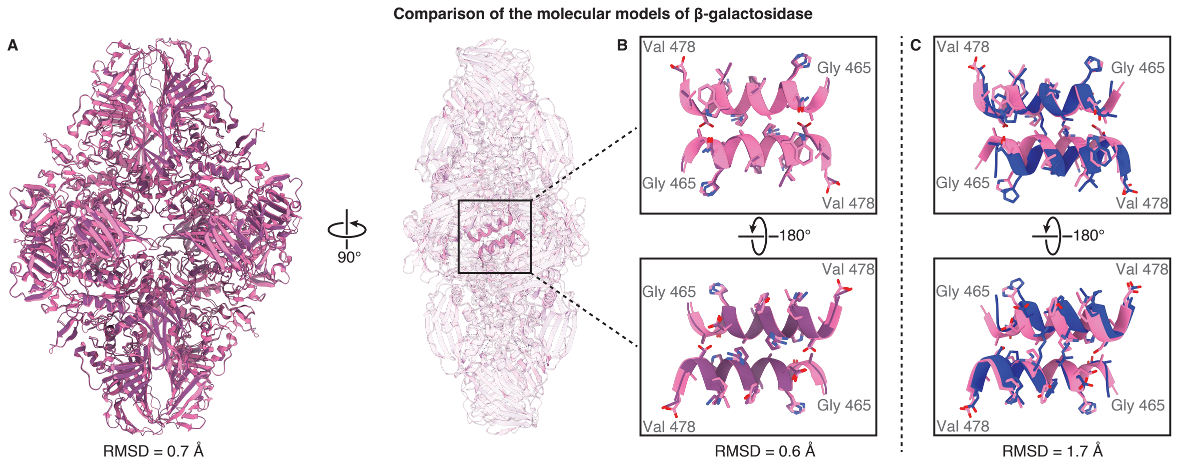


**Figure S12 | Comparison of the molecular models for soft-landed, rehydrated, and native β-galactosidase.** (**A**) Comparison of the models for the rehydrated (best) and native structures (purple and pink, respectively). (**B**) Comparison of the subunit contacts within the well resolved core between the rehydrated (best) and native structures. The models closely resemble each other, with an RMSD of 0.6 Å. (**C**) Comparison of the same subunit contacts in the dehydrated (soft-landed with ice, PDB: 8Q7Y[3]) and native structures (blue and pink, respectively). The models show larger differences with an RMSD of 1.7 Å.

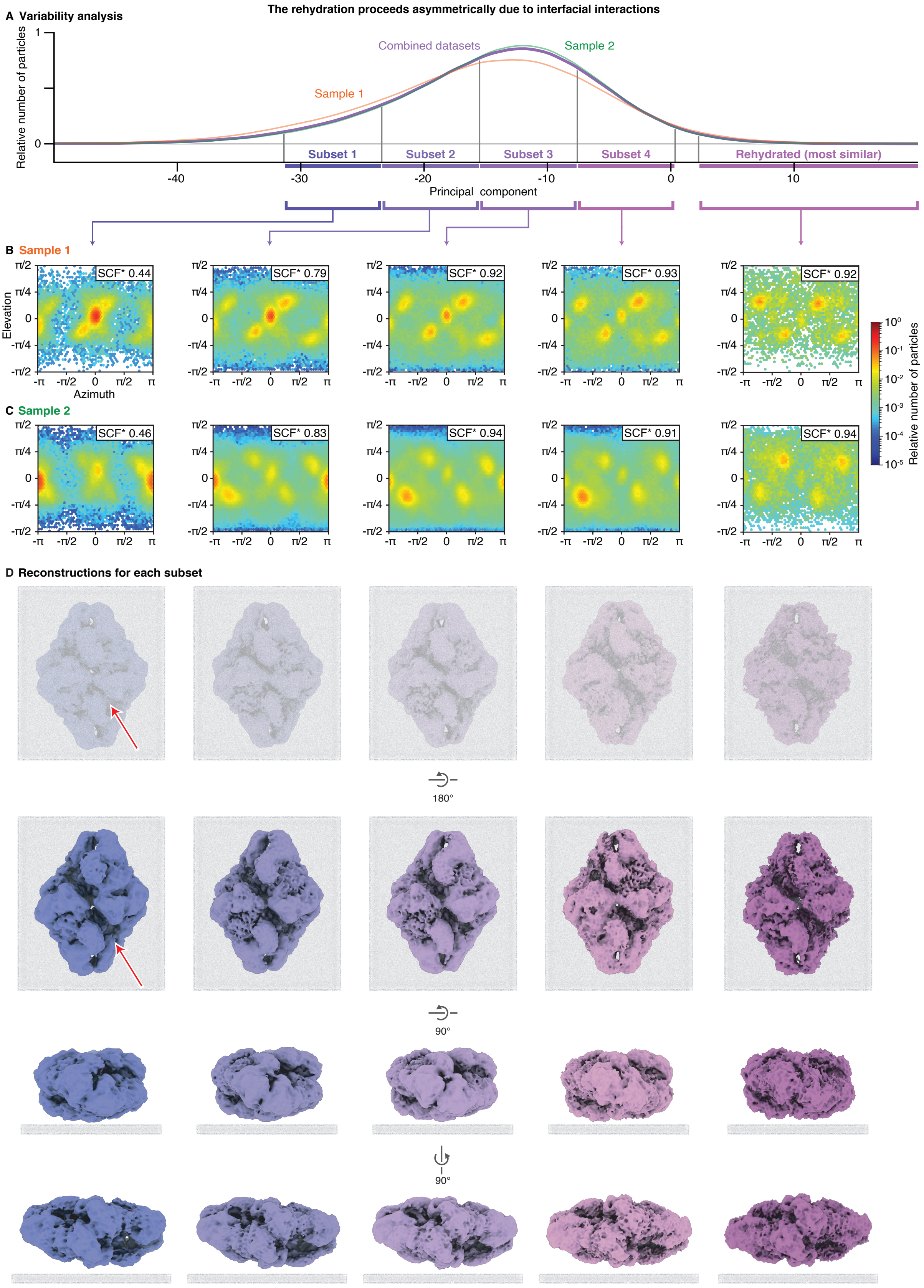


**Figure S13 | The rehydration proceeds asymmetrically due to interfacial interactions.** (**A**) Distribution of the rehydrated β-galactosidase particles along the first variability component for the entire dataset collected from two different samples (purple, from Fig. 4D), as well as for each of the samples (orange and green). The central portion of the distribution is subdivided into 4 subsets. The subset of particles most similar to the native state that are included in the reconstruction in Fig. 3D is also indicated. (**B**) Angular distributions of the particles in each subset for the first sample, with the corresponding SCF* values indicated (from Fig. 4F–I). Along the principal component, preferred orientation decreases. (**C**) Angular distributions for the second sample. During imaging, this sample was apparently mounted with the opposite side facing up compared to the first sample, which shifts the angular distribution by 180 degrees. This can for example be seen for the most asymmetric particles (subset 1), whose preferred orientation (adsorption of the particles with one of their faces) is rotated by 180 degrees. Note that besides this rotation, the angular distributions for sample 2 also show other changes to the preferred orientations of the particles, which is likely the result of small differences in the interfacial properties of the two samples. (**D**) Reconstructions for each subset for the combined particles of both samples. The map of the first subset shows significant asymmetry, with one face of the protein more expanded than the other. Arrows highlight the asymmetry in the subunit contacts on both faces. As preferred orientation decreases along the principal component, the volumes become more symmetric. This suggests that the interfacial interactions that lead to preferred orientation also hinder the rehydration of the particles, causing them to become asymmetric. The putative interaction with the interface is schematically indicated (grey). All reconstructions are performed with $C_1$ symmetry. The maps are displayed with thresholds of 5 σ above the mean.

#### J. Supplementary Tables

**Table S1 | Data collection parameters as well as reconstruction and modelling statistics for GroEL.**

| Figure | Fig 2D | Fig 2A | Fig 2B | Fig 2C |
|---|---|---|---|---|
| **Name** | **Plunge-frozen** | **Soft-landed without ice** | **Soft-landed with ice** | **Soft-landed and rehydrated** |
| EMDB accession | EMD-52282 | EMD-52627 | EMD-52626 | EMD-52284 |
| **Data collection** | | | | |
| Microscope | Thermo Fisher Titan Krios G4 | Thermo Fisher Talos Arctica | Thermo Fisher Titan Krios G3 | Thermo Fisher Titan Krios G4 |
| Detector | Falcon 4i | Falcon 4i | Gatan K3 | Falcon 4i |
| Energy Filter | SelectrisX | - | GIF Bioquantum | SelectrixX |
| Energy Slit | 10 eV | - | 20 eV | 10 eV |
| Voltage (kV) | 300 | 200 | 300 | 300 |
| Electron exposure ($e^{-}/Å^2$) | 60 | 40 | 40 | 60 |
| Magnification | 165,000 | 105,000 | 105,000 | 165,000 |
| Defocus range (μm) | -0.8 – -2.2 | -1.0 – -2.25 | -1.0 – -2.5 | -0.8 – -2.2 |
| Pixel size (Å) | 0.726 | 1.17 | 0.83 | 0.726 |
| **Data processing** | | | | |
| Initial movies (no.) | 9,759 | 111 | 4,004 | 44,000 |
| Symmetry imposed | $D_7$ | $D_7$ | $D_7$ | $D_7$ |
| Initial particle images (no.) | 1,894,809 | 47,428 | 904,984 | 3,455,984 |
| Final particle images (no.) | 366,916 | 26,085 | 75,264 | 119,637 |
| Map resolution (Å) | 1.9 | 6.7 | 4.8 | 3.9 |
| FSC threshold | 0.143 | 0.143 | 0.143 | 0.143 |
| Map resolution range (Å) | 1.6 – 4.0 | 6.2 – 10 | 4.5 - 9.4 | 3.5 – 12.0 |

**Table S2 | Data collection parameters as well as reconstruction and modelling statistics for β-galactosidase.**

| Figure | Fig 3E | Fig 3A | Fig 3C | Fig 3D |
|---|---|---|---|---|
| **Name** | **Plunge-frozen** | **Soft-landed without ice** | **Soft-landed and rehydrated (Average)** | **Soft-landed and rehydrated (Best)** |
| EMDB accession | EMD-52230 | EMD-52274 | EMD-52244 | EMD-52260 |
| PDB accession | 9HKI | | | 9HLM |
| **Data collection** | | | | |
| Microscope | Thermo Fisher Titan Krios G4 | | | |
| Detector | Falcon 4i | | | |
| Energy Filter | SelectrisX | | | |
| Energy Slit | 10 eV | | | |
| Voltage (kV) | 300 | | | |
| Electron exposure ($e^-/Å^2$) | 60 | | | |
| Magnification | 165,000 | 130,000 | 165,000 | 165,000 |
| Defocus range (μm) | -0.8 – -2.2 | | | |
| Pixel size (Å) | 0.726 | 0.926 | 0.726 | 0.726 |
| **Data processing** | | | | |
| Initial movies (no.) | 11,618 | 945 | 44,929 | 44,929 |
| Symmetry imposed | $D_2$ | $D_2$ | $C_1$ | $D_2$ |
| Initial particle images (no.) | 1,505,585 | 104,418 | 4,415,932 | 4,415,932 |
| Final particle images (no.) | 305,360 | 65,983 | 1,504,908 | 25,206 |
| Map resolution (Å) | 1.9 | 5.9 | 2.9 | 3.7 |
| FSC threshold | 0.143 | 0.143 | 0.143 | 0.143 |
| Map resolution range (Å) | 1.6 – 3.5 | 4.0 – 12.6 | 1.6 – 5.3 | 3.2 – 9.0 |
| **Refinement** | | No atomic model built | No atomic model built | |
| Initial model used (PDB ID) | 6CVM | - | - | 6CVM |
| Model resolution (Å) | 1.8 | - | - | 3.8 |
| FSC threshold | 0.143 | - | - | 0.143 |
| Model resolution range (Å) | 1.8/1.8/1.9 | - | - | 3.6/3.8/3.9 |
| FSC threshold | 0/0.143/0.5 | - | - | 0/0.143/0.5 |
| Map sharpening *B* factor ($Å^2$) | 41.9 | - | - | 62.7 |
| Model composition | | | | |
| Non-hydrogen atoms | 36752 | - | - | 32040 |
| Protein residues | 4084 | - | - | 3984 |

| | | | | |
|---|---|---|---|---|
| Waters | 3952 | - | - | - |
| Ligands | 12 | - | - | - |
| Average *B* factors (Å$^2$) | | | | |
| Protein | 10.02 | - | - | 96.18 |
| Ligand | 8.52 | - | - | - |
| Water | 10.80 | | | - |
| R.m.s. deviations | | | | |
| Bond lengths (Å) | 0.004 | - | - | 0.005 |
| Bond angles (°) | 1.036 | - | - | 1.026 |
| Validation | | | | |
| MolProbity score | 1.08 | - | - | 2.17 |
| Clashscore | 2.74 | - | - | 21.26 |
| Poor rotamers (%) | 0.34 | - | - | 0.23 |
| Ramachandran plot | | | | |
| Favored (%) | 97.94 | - | - | 95.03 |
| Allowed (%) | 2.06 | - | - | 4.67 |
| Disallowed (%) | 0.00 | - | - | 0.30 |
